# The Association between *SLC25A15* Gene Polymorphisms and Hyperornithinemia-hyperammonemia-homocitrullinuria Syndrome: Using In Silico Analysis

**DOI:** 10.1101/786301

**Authors:** Nuha A. Mahmoud, Dina T. Ahmed, Zainab O. Mohammed, Fatima A. Altyeb, Mujahed I. Mustafa, Mohamed A. Hassan

## Abstract

**Background:** Hyperornithinemia-hyperammonemia-homocitrullinuria (HHH) syndrome is an autosomal recessive inborn error of the urea cycle. It is caused by mutations in the *SLC25A15* gene that codes the mitochondrial ornithine transporter. The aim of this study is to detect and identify the pathogenic SNPs in *SLC25A15* gene through a combination set of bioinformatics tools and their effect on the structure and function of the protein.

**Methods:** The deleterious SNPs in *SLC25A15* are detected by various bioinformatics tools, with addition to identifying their effects on the structure and function of this gene.

**Results:** 20 deleterious SNPs out 287of were found to have their own damaging effects on the structure and function of the *SLC25A15* gene.

**Conclusion:** This study is the first in silico analysis of SLC25A15 using a selection of bioinformatics tools to detect functional and structural effects of deleterious SNPs. Finding the pathogenic SNPs is a promising start to innovate new, useful SNP diagnostic markers for medical testing and for safer novel therapies specifically targeting mutant *SLC25A15*.

## 1. Introduction

Hyperornithinemia-hyperammonemia-homocitrullinuria (HHH) syndrome is an autosomal recessive inborn error of the urea cycle (1–5), (OMIM#238970) (6–8), it is located on chromosome 13q14(7, 9). This syndrome occurs in the early infancy or childhood period, although, few cases of adult onset have also been reported (7, 10–14). HHH syndrome is caused by mutations in the *SLC25A15* gene that codes the mitochondrial ornithine transporter, which transports ornithine across the inner mitochondrial membrane from the cytosol to the mitochondrial matrix (15). This is a vital step in the urea cycle, which serves to eliminate toxic ammonium ions from the breakdown of nitrogen (6, 16–21). The HHH clinical symptoms are highly variable, which include spastic paraplegia, extrapyramidal signs, stroke-like episodes, hypotonia, seizures, ataxia, protein intolerance, failure to thrive, pyramidal dysfunction developmental delay/intellectual disability, hepatic failure (13, 16, 20, 22–24), and even death can arise if the condition is not diagnosed early and treated quickly (25).

Even though, this disease targets a wide variation of different ethnicities, Canada (23%), Italy (17%) and Japan (13%) were reported with the highest prevalence rates (15, 26–28)

Albeit HHH treatment includes a strict treatment protocol consisting control of dietary protein, nitrogen forager medications, as well as, amino acid and vitamins supplementations, still there is no effective therapy for this type of inborn disease (4, 22–25), thus identifying the *SLC25A15* gene holds a promising future tool to improve both survival rates as well as the quality of life in surviving of HHH patients(29, 30).

Detection and identification of pathogenic SNPs of *SLC25A15* through different in silico prediction software are the main aims of this study. As well as, to determine the structure, bio functions and the regulatory mechanism of their respective proteins. This study is the first in silico analysis of the SLC25A15 coding region. Using this approach to identify the pathogenic SNPs with no cost; novel therapies specifically targeting mutant *SLC25A15* will open new methods of treatment for HHH patients(31).

## 2. Methods

### 2.1 Data Mining

The *SLC25A15* human gene data was obtained from the National Center for Biotechnology Information (NCBI) (http://www.ncbi.nlm.nih.gov/). The variant’s information of the gene was collected from the NCBI dbSNP database (http://www.ncbi.nlm.nih.gov/snp/) and the SLC25A15 protein sequence was obtained from UniProt database (https://www.uniprot.org/).

### 2.2 SIFT

The Sorting Intolerant from Tolerant (SIFT) algorithm is a standardized tool for detecting all missense SNPs in the protein sequence by searching for homologous sequences and carry out multiple sequence alignment. The score of each variant ranges from 0 to 1, deleterious SNP is considered when the score is ≤0.05, while tolerated SNP’s score is >0.05 (32, 33) SIFT is an available online tool (34) (http://sift-dna.org).

### 2.3 PolyPhen-2

PolyPhen-2 (polymorphism phenotyping v2) is a tool that predicts the impact of substitution of the amino acid on the structure and function of the protein. The input data needs accession number, position of variants, original and mutant amino acids. The PolyPhen figures the PSIC (position-specific independent count) and counts the score difference between the two variants. When resulted with a score of (0.96–1) it’s considered ‘Probably damaging’, while (0.71–0.95) scores considered as ‘Possibly damaging and Benign’ (0.31–0.7) (32, 35, 36). PolyPhen-2 is available at (http://genetics.bwh.harvard.edu/pph2/).

### 2.4 PROVEAN

PROVEAN (Protein Variation Effect Analyzer), is an algorithm, which accurately predicts the impact of single amino acid substitutions, multiple substitutions, insertions and deletions(37). The tool accepts a protein sequence and amino acid variations as input, performs a BLAST to identify homologous sequences, and generates the scores. A variant is predicted to be ‘deleterious’ if the final score is below −2.5, and is predicted to be ‘neutral’ otherwise (37, 38). It is available at (http://provean.jcvi.org/index.php).

### 2.5 PHD-SNP

Predictor of Human Deleterious Single Nucleotide Polymorphisms; is an online Support Vector Machine SVM-based classifier. It predicts whether the amino acid substitution is deleterious or neutral (39). It is available at (http://snps.biofold.org/phd-snp/phd-snp.html).

### 2.6 SNPs&GO

Single Nucleotide Polymorphism Database & Gene Ontology is a support vector machine (SVM) based method. The information derived from the protein sequence, structure, and protein function; it needs a whole protein in FASTA sequence, and/or its three-dimensional structure, a target SNP and its functional Gene Ontology (GO) terms to be an input option. The output provides, for each protein variation, the probabilities to be disease or neutral. The score of predicted variants is 82% accurate with Matthews’s correlation coefficient of 0.63 and if it is > 0.5 this is defined as a ‘disease’ (39–41). It is freely available at (http://snps.biofold.org/snps-and-go/snps-and-go.html)

### 2.7 PANTHER

Protein Analysis through evolutionary relationships, is used for analysis of variants, it utilizes hidden Markov model (HMM) based statistical methods and multiple sequence alignments which create the substitution position-specific evolutionary conservation score (subPSEC). The SNPs is considered deleterious with score (< −3) and neutral with (score > −3) (39). It is available at (http://www.pantherdb.org/)

### 2.8 P-Mut

P-Mut is an online tool annotates and predicts the damaging mutations. Pmut is rely on sequence information of any protein to identify the mutations. It delivers a very simple output: a yes/no answer and a reliability index. Its methods performance is 84 % overall success rate, and 67 % improvement over random (42–44). It is available at (http://mmb.irbbarcelona.org/PMut/).

### 2.9 I-Mutant 3.0

I-Mutant is a neural-network-based tool for prediction of protein stability and modulation upon single amino acid substitution. The protein sequence and variant position is needed for input data. The output is the predicted free energy change (DDG), which classifies the prediction into: ‘large decrease’ (DDG < −0.5 kcal/mol), ‘large increase’ (DDG > 0.5 kcal/mol), or ‘neutral’ (−0.5 < DDG < 0.5 kcal/mol) (39, 44). It is available at (http://gpcr2.biocomp.unibo.it/cgi/predictors/I-Mutant3.0/I-Mutant3.0.cgi).

### 2.10 MUpro

A support vector machine-based bioinformatics tool, MUpro, is used for predicting the stability changes of a protein upon nonsynonymous SNPs. The value of the energy change is assumed, and a confidence score among −1 and 1 for measuring the confidence of the assumption is calculated. A score <0 shows that the variable reduces the protein stabilization and vice versa (45). It is available at (http://mupro.proteomics.ics.uci.edu/).

### 2.12 Project HOPE

Have Our Protein Explained, is an online next-generation tool used for mutant analysis. HOPE’s layout demonstrates the molecular origin of disease’s phenotype which results from protein mutation. The Web services and DAS servers are used to collect the data. The sources from which HOPE gathers the data are ‘Protein’s 3D structure’ and the ‘UniProt database’, of well-annotated protein sequences. For each protein, this data is stored in a PostgreSQL-based information system. The data is processed and used to predict the mutation impact on the 3D structure and the function of the protein. The report is generated, is supported with figures and clarify the effects of mutation. Lastly the report can be utilized to design follow-up experiments, better diagnostic & medicines (46). Available at (http://www.cmbi.ru.nl/hope/input/).

### 2.11 RaptorX

Since the human *SLC25A15* protein’s 3D structure is unavailable in the Protein Data Bank, RaptorX software was used to obtain the 3D structural model for wild-type of SLC25A15. RaptorX is a bioinformatics web server predicts the structural property of a protein sequence without the need of any templates. It surpasses other servers, chiefly for proteins without close homologs in PDB or with very sparse sequence profile (47). It is available at (http://raptorx.uchicago.edu/).

### 2.12 UCSF Chimera

This desktop application is for the interactive visualization and analysis of molecular structures and related data. The application features are expanded due to incorporation of the web access with desktop tool. The features include the accomplishing computationally intense work and is able to connect to databases. Chimera is subdivided into a ‘Core’, which is responsible for basic services and visualization, and ‘Extensions’, that contain new features with higher functionality.

These extensions are Multialign Viewer, docking, trajectories, visualize large-scale molecular assemblies, etc. chimera images with high resolution. This application can generate an animation (48). Chimera is available at (http://www.cgl.ucsf.edu/chimera/).

### 2.13 GeneMANIA

GeneMANIA is an online server generating proposition about gene function, investigating gene lists and prioritizing genes for functional assays. It expands the query list with functionally similar genes that recognizes by using available genomics and proteomics data. The high accuracy of the GeneMANIA prediction algorithm and large database make the GeneMANIA an useful tool for any biologist (49, 50). It is available at (http://genemania.org/).

## 3. Result and discussion

### 3.1 Result

### 3.2 Discussion

Various bioinformatics tools were performed to identify the SNPs impact in the *SLC25A15* protein. 20 mutations have been distinguished to affect the stability and function of SLC25A15, 12 out of 20 are considered as novel (D31H, D31Y, Y64C, G66S, G86C, G168E, T176A, F188S, K234I, Y287H, E288K and E288G). To ensure the pathological effect of the mutations at the molecular level, different parameters plus trustworthy information were used, and for more accuracy, numerous methods rather than one method (PolyPhen-2, PROVEAN, PHD-SNP, SNP&GO, PANTHER, P-MUT, I-Mutant3.0, MUpro, Project HOPE, RaptorX, Chimera, and GeneMANIA were used (Figure1)

**Figure 1:**
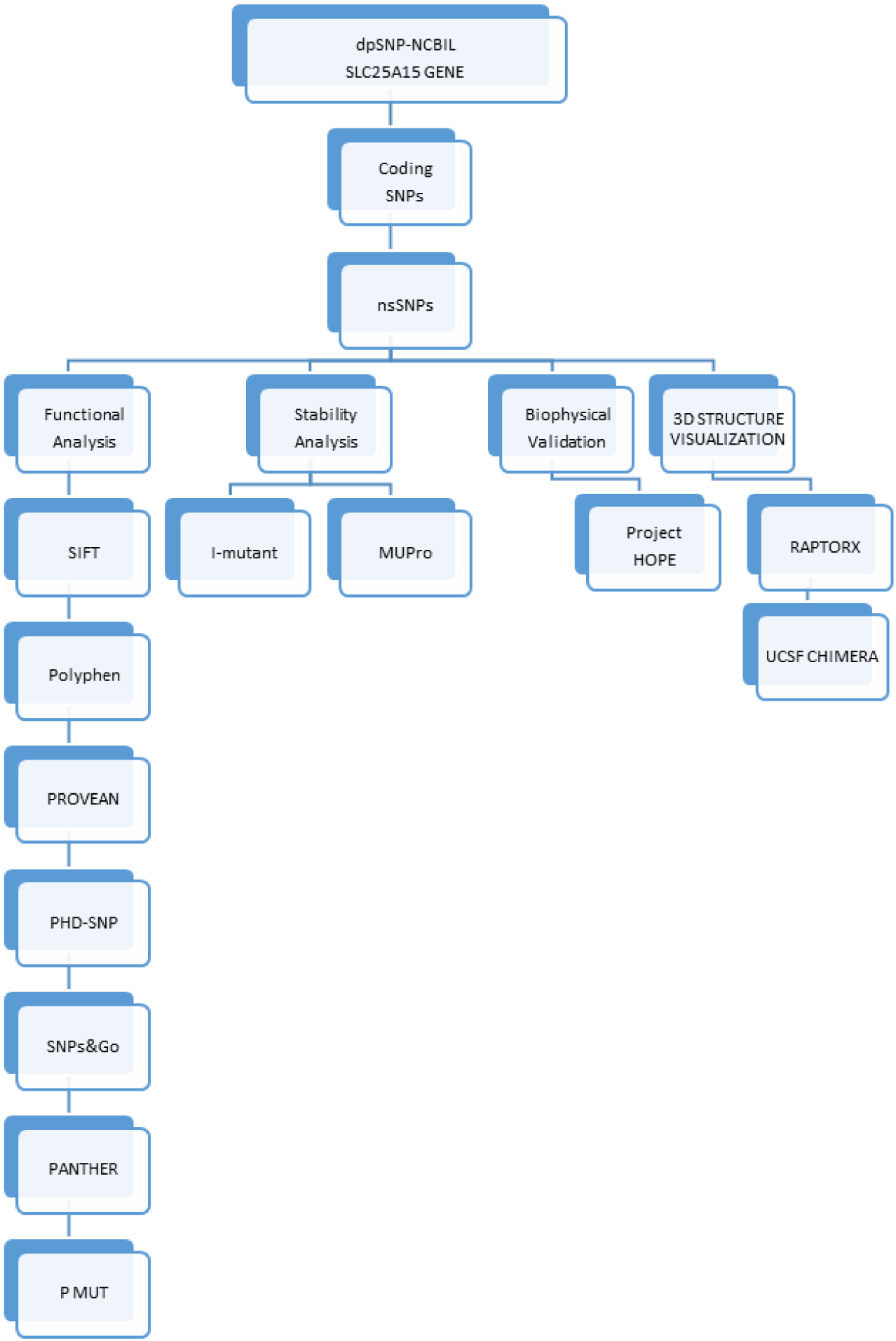
Diagrammatic representation of *SLC25A15* gene in silico work flow.

630 variants of the Homo Sapiens *SLC25A15* gene which situated on the coding region were acquired from dbSNP/NCBI Database. To predict if these variants affect the function of the protein, phenotype and end by disease 220 missense mutations were submitted to SIFT, PolyPhen-2 and PROVEAN software respectively. 57 SNPs were predicted to be ‘deleterious’ by SIFT software, while by PolypPhen-2; 114 SNPs were predicted to be ‘damaging’, in which 79 were ‘probably damaging’ and 35 as ‘possibly damaging ‘, and PROVEAN predicted 142 SNPs to be deleterious. The triple positive results of these software were 27 SNPs, and they are affecting the function of the protein (Table 1). These 27 pathogenic SNPs were submitted to PHD-SNP, SNP&GO, PANTHER and P-MUT for further confirmation; 27 deleterious SNPs were predicted by PHD-SNP and SNP&GO while PANTHER predict 25 variants and P-MUT predicted 21 deleterious SNPs (table 2 AND 3). The fourth positive results of these software were 20 pathogenic SNPs, were submitted to I-Mutant and MUpro servers in order to examine the effect of mutation on the stability of the protein. I-Mutant revealed all these 20 pathogenic SNPs decreased the stability of the protein except for 4 SNPs with I-Mutant (D31Y, G168E, G216S, and G217R), but one SNP with MUPRO (K234I) was predicted to increase the stability (Table 4).

**Table (1):** Deleterious – damaging (Affected) nsSNPs predicted by various software:

| Amino acid change | SIFT |  | POLYPHEN-2 |  | PROVEAN |  |
| --- | --- | --- | --- | --- | --- | --- |
|  | Prediction | Score | Prediction | Score | Prediction (cutoff=-2.5) | Score |
| D31H | AFFECTED | 0.00 | probably damaging | 1 | Deleterious | -6.651 |
| D31Y | AFFECTED | 0.00 | probably damaging | 1 | Deleterious | -8.551 |
| Y64C | AFFECTED | 0.00 | probably damaging | 1 | Deleterious | -8.418 |
| G66S | AFFECTED | 0.00 | probably damaging | 1 | Deleterious | -5.634 |
| L71Q | AFFECTED | 0.00 | probably damaging | 1 | Deleterious | -4.892 |
| G86C | AFFECTED | 0.00 | probably damaging | 1 | Deleterious | -8.585 |
| G113C | AFFECTED | 0.00 | probably damaging | 1 | Deleterious | -8.512 |
| G168E | AFFECTED | 0.00 | probably damaging | 1 | Deleterious | -6.745 |
| T176A | AFFECTED | 0.00 | probably damaging | 1 | Deleterious | -4.81 |
| F188S | AFFECTED | 0.00 | probably damaging | 1 | Deleterious | -7.651 |
| G216S | AFFECTED | 0.00 | probably damaging | 1 | Deleterious | -5.543 |
| G217R | AFFECTED | 0.00 | probably damaging | 1 | Deleterious | -5.213 |
| G220R | AFFECTED | 0.00 | probably damaging | 1 | Deleterious | -7.39 |
| K234I | AFFECTED | 0.00 | probably damaging | 1 | Deleterious | -7.488 |
| Q238R | AFFECTED | 0.00 | probably damaging | 1 | Deleterious | -3.795 |
| R275Q | AFFECTED | 0.00 | probably damaging | 1 | Deleterious | -3.64 |
| R275L | AFFECTED | 0.00 | probably damaging | 1 | Deleterious | -6.303 |
| Y287H | AFFECTED | 0.00 | probably damaging | 1 | Deleterious | -4.617 |
| E288K | AFFECTED | 0.00 | probably damaging | 1 | Deleterious | -3.773 |
| E288G | AFFECTED | 0.00 | probably damaging | 1 | Deleterious | -6.603 |

**Table (2):** Disease-associated nsSNPs predicted by various software:

| Amino acid change | SNP&GO |  |  | PHD-SNP |  |  | PANTHER |  |  |
| --- | --- | --- | --- | --- | --- | --- | --- | --- | --- |
|  | Prediction | RI | Score | Prediction | RI | Score | Prediction | RI | Score |
| D31H | Disease | 4 | 0.719 | Disease | 7 | 0.863 | Disease | 10 | 0.992! |
| D31Y | Disease | 7 | 0.842 | Disease | 9 | 0.933 | Disease | 10 | 0.995 |
| Y64C | Disease | 6 | 0.797 | Disease | 8 | 0.915 | Disease | 8 | 0.905 |
| G66S | Disease | 6 | 0.794 | Disease | 8 | 0.908 | Disease | 10 | 1 |
| L71Q | Disease | 5 | 0.77 | Disease | 6 | 0.793 | Disease | 6 | 0.82 |
| G86C | Disease | 1 | 0.539 | Disease | 4 | 0.721 | Disease | 5 | 0.775 |
| G113C | Disease | 6 | 0.801 | Disease | 8 | 0.89 | Disease | 8 | 0.916 |
| G168E | Disease | 7 | 0.826 | Disease | 7 | 0.828 | Disease | 7 | 0.85 |
| T176A | Disease | 2 | 0.609 | Disease | 1 | 0.574 | Disease | 0 | 0.525 |
| F188S | Disease | 6 | 0.78 | Disease | 8 | 0.877 | Disease | 6 | 0.814 |
| G216S | Disease | 4 | 0.718 | Disease | 6 | 0.793 | Disease | 10 | 0.988 |
| G217R | Disease | 6 | 0.779 | Disease | 7 | 0.874 | Disease | 2 | 0.618 |
| G220R | Disease | 7 | 0.841 | Disease | 8 | 0.914 | Disease | 8 | 0.882 |
| K234I | Disease | 6 | 0.799 | Disease | 7 | 0.874 | Disease | 10 | 0.999 |
| Q238R | Disease | 4 | 0.7 | Disease | 6 | 0.799 | Disease | 3 | 0.633 |
| R275Q | Disease | 5 | 0.76 | Disease | 7 | 0.847 | Disease | 10 | 0.986 |
| R275L | Disease | 7 | 0.851 | Disease | 9 | 0.932 | Disease | 10 | 0.991 |
| Y287H | Disease | 3 | 0.649 | Disease | 7 | 0.868 | Disease | 3 | 0.65 |
| E288K | Disease | 5 | 0.743 | Disease | 9 | 0.943 | Disease | 0 | 0.503 |
| E288G | Disease | 1 | 0.537 | Disease | 7 | 0.875 | Disease | 1 | 0.558 |

**Table (3):** Disease-associated nsSNPs predicted by P-Mut software:

| Amino acid change | P-Mut Prediction | P-Mut Score |
| --- | --- | --- |
| D31H | Disease | 0.85 (91%) |
| D31Y | Disease | 0.86 (91%) |
| Y64C | Disease | 0.85 (91%) |
| G66S | Disease | 0.85 (91%) |
| L71Q | Disease | 0.76 (88%) |
| G86C | Disease | 0.76 (88%) |
| G113C | Disease | 0.86 (91%) |
| G168E | Disease | 0.73 (87%) |
| T176A | Disease | 0.71 (86%) |
| F188S | Disease | 0.86 (91%) |
| G216S | Disease | 0.63 (83%) |
| G217R | Disease | 0.86 (91%) |
| G220R | Disease | 0.86 (91%) |
| K234I | Disease | 0.86 (91%) |
| Q238R | Disease | 0.74 (87%) |
| R275Q | Disease | 0.86 (91%) |
| R275L | Disease | 0.86 (91%) |
| Y287H | Disease | 0.80 (89%) |
| E288K | Disease | 0.85 (91%) |
| E288G | Disease | 0.85 (91%) |

**Table (4):** The effect of the deleterious SNPs on the stability predicted by 2 software:

| Amino acid change | I- mutant |  |  | MUPRO |  |
| --- | --- | --- | --- | --- | --- |
|  | SVM2 Prediction Effect | RI | DDG Value Prediction | Prediction | Score |
| D31H | Decrease | -0.16 | 1 | Decrease | -1.1775469 |
| D31Y | Increase | 0.46 | 3 | Decrease | -0.68362433 |
| Y64C | Decrease | -1.33 | 3 | Decrease | 1.09234 |
| G66S | Decrease | -1.5 | 9 | Decrease | -0.66844573 |
| L71Q | Decrease | -2.24 | 9 | Decrease | -1.8046296 |
| G86C | Decrease | 6 | -1.12 | Decrease | -0.46574415 |
| G113C | Decrease | 1 | -0.64 | Decrease | -0.13197126 |
| G168E | Increase | 3 | -0.54 | Decrease | -0.40969468 |
| T176A | Decrease | 8 | -1.18 | Decrease | -1.526078 |
| F188S | Decrease | 7 | -1.62 | Decrease | -1.740386 |
| G216S | Increase | -0.76 | 1 | Decrease | -0.98156517 |
| G217R | Increase | -0.34 | 1 | Decrease | -1.1923443 |
| G220R | Decrease | -0.39 | 1 | Decrease | -0.50470067 |
| K234I | Decrease | 0.19 | 1 | Increase | 0.014638836 |
| Q238R | Decrease | -0.22 | 2 | Decrease | -1.0081246 |
| R275Q | Decrease | -0.93 | 7 | Decrease | -0.6423489 |
| R275L | Decrease | -0.32 | 7 | Decrease | -0.007036716 |
| Y287H | Decrease | -1.37 | 6 | Decrease | -1.6533478 |
| E288K | Decrease | -0.75 | 8 | Decrease | -1.1400911 |
| E288G | Decrease | -1.14 | 7 | Decrease | -1.3861129 |

Project HOPE reveled that some mutant residues (which are buried in the core of the protein) are larger than the wild-type residues, this makes the new residue in an improper position to make the same hydrogen bond as the original wild-type residue did, while the smaller sized mutants cannot fill the core of the protein. It also showed the mutation may change the charge of the residue and lead to repulsion of ligands or other residues with the same charge, there are 9 variants remain unchanged. In Project HOPE results, 12 variants with mutant residue are more hydrophobic than the wild-type residue, these hydrophobicity variations can affect the hydrophobic interactions with the membrane lipids, and it identified 10 variants with transmembrane domain as well. The amino acid Glycine is the most flexible of all residues, this flexibility might be necessary for the protein’s function. Thus mutation of this glycine can abolish this function. For visualization of the wild and mutant types of amino acids in SLC25A15, Chimera software was performed (Figures 2–21)

**Figure 2:**
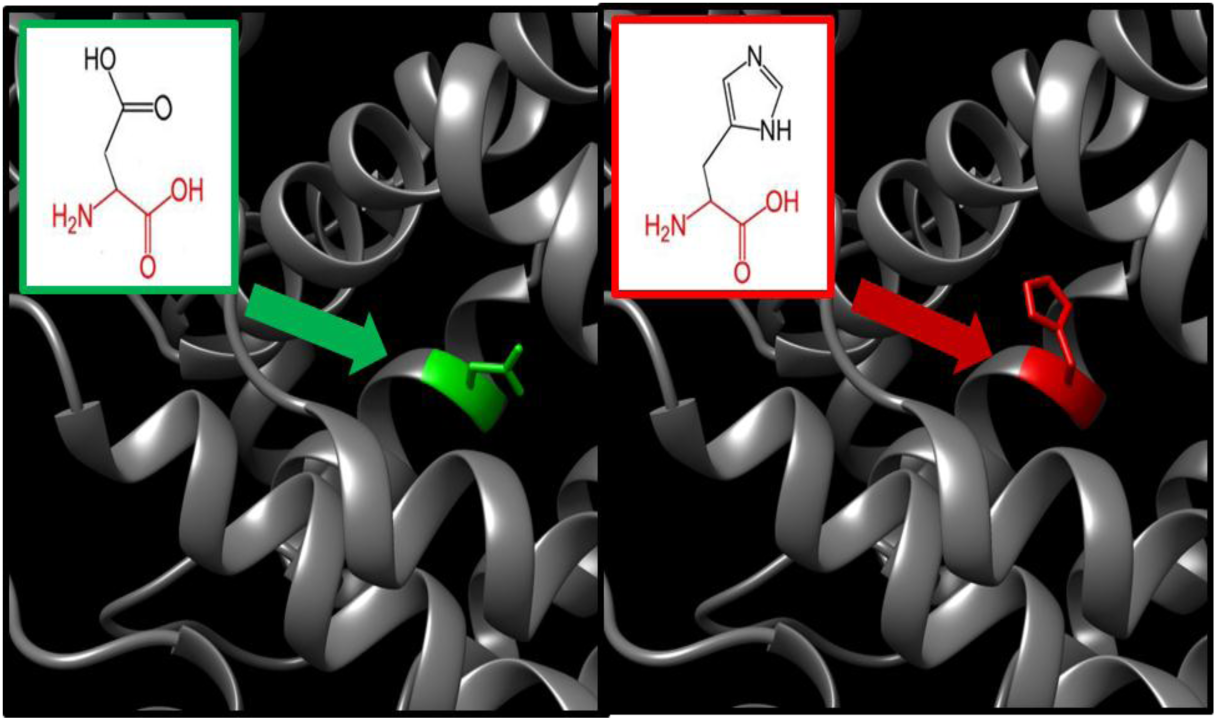
(G31H) the amino acid Aspartic Acid (green) changes to Histidine (red) at position 31.

**Figure 3:**
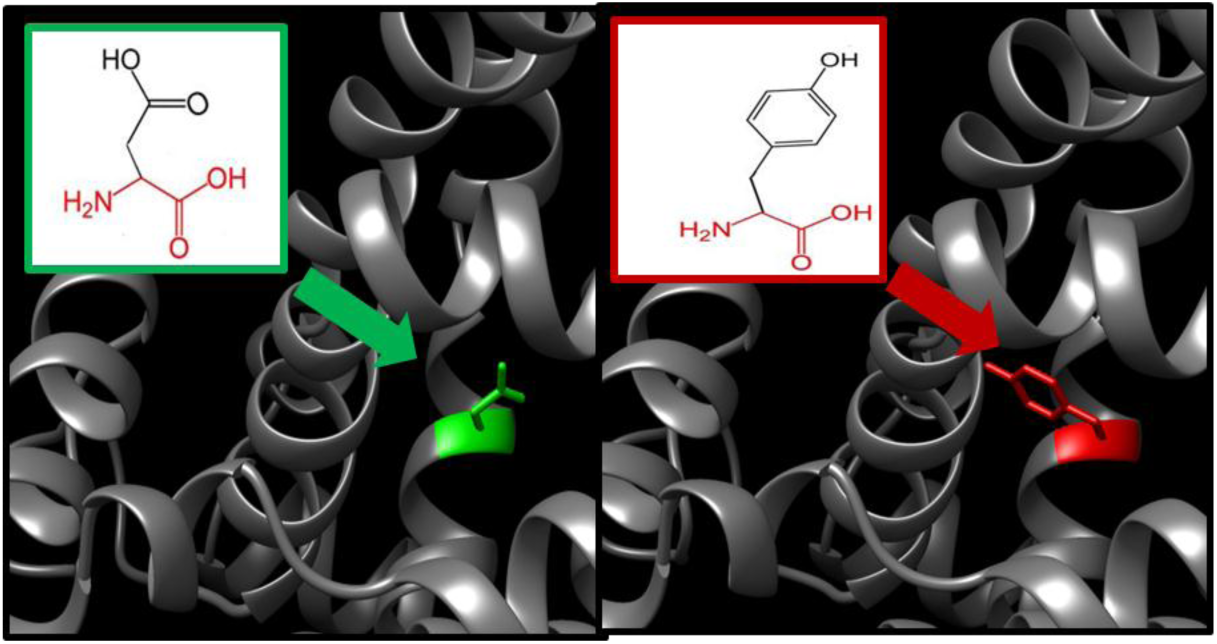
(D31Y) the amino acid Aspartic Acid (green) changes to Tyrosine (red) at position 31.

**Figure 4:**
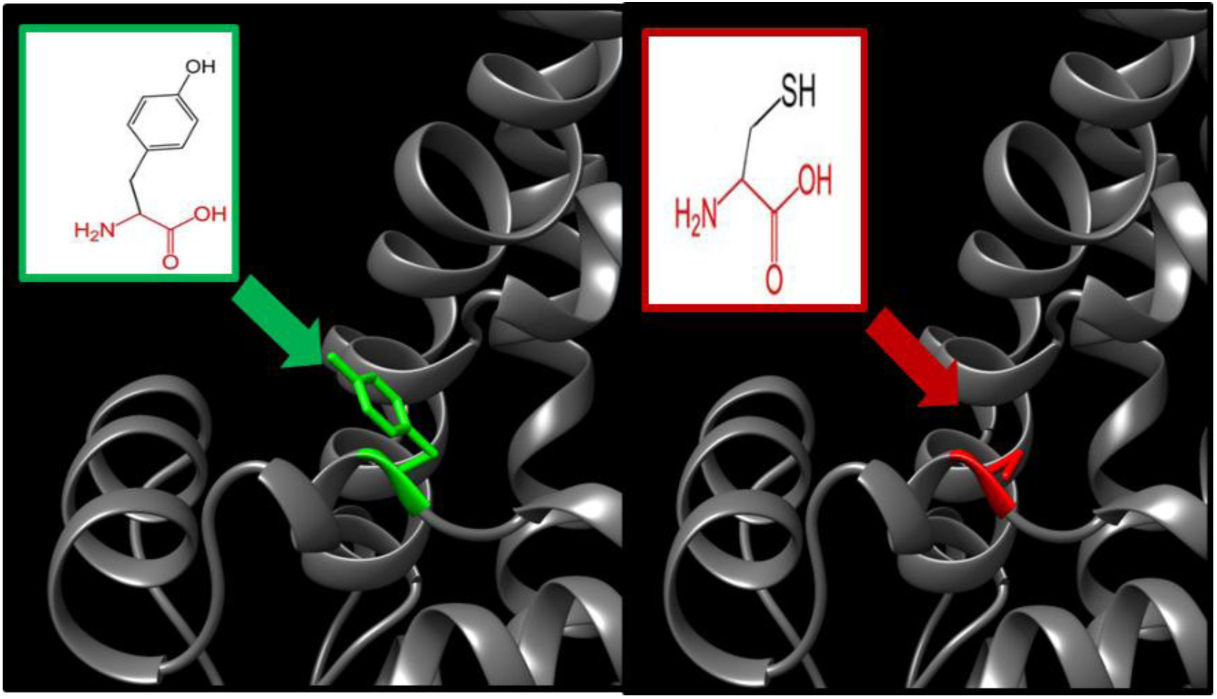
(Y64C) the amino acid Tyrosine (green) changes into a Cysteine (red) at position 64

**Figure 5:**
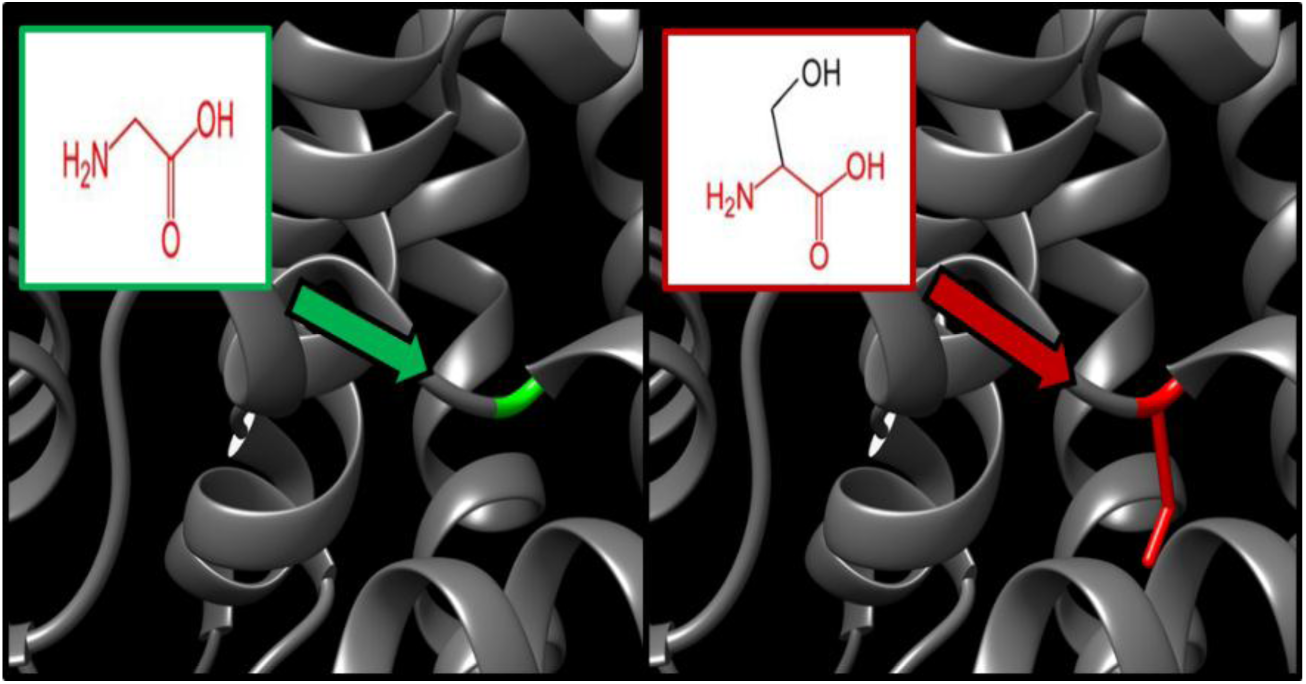
(G66S) the amino acid Glycine (green) changes into a Serine (red) at position 66.

**Figure 6:**
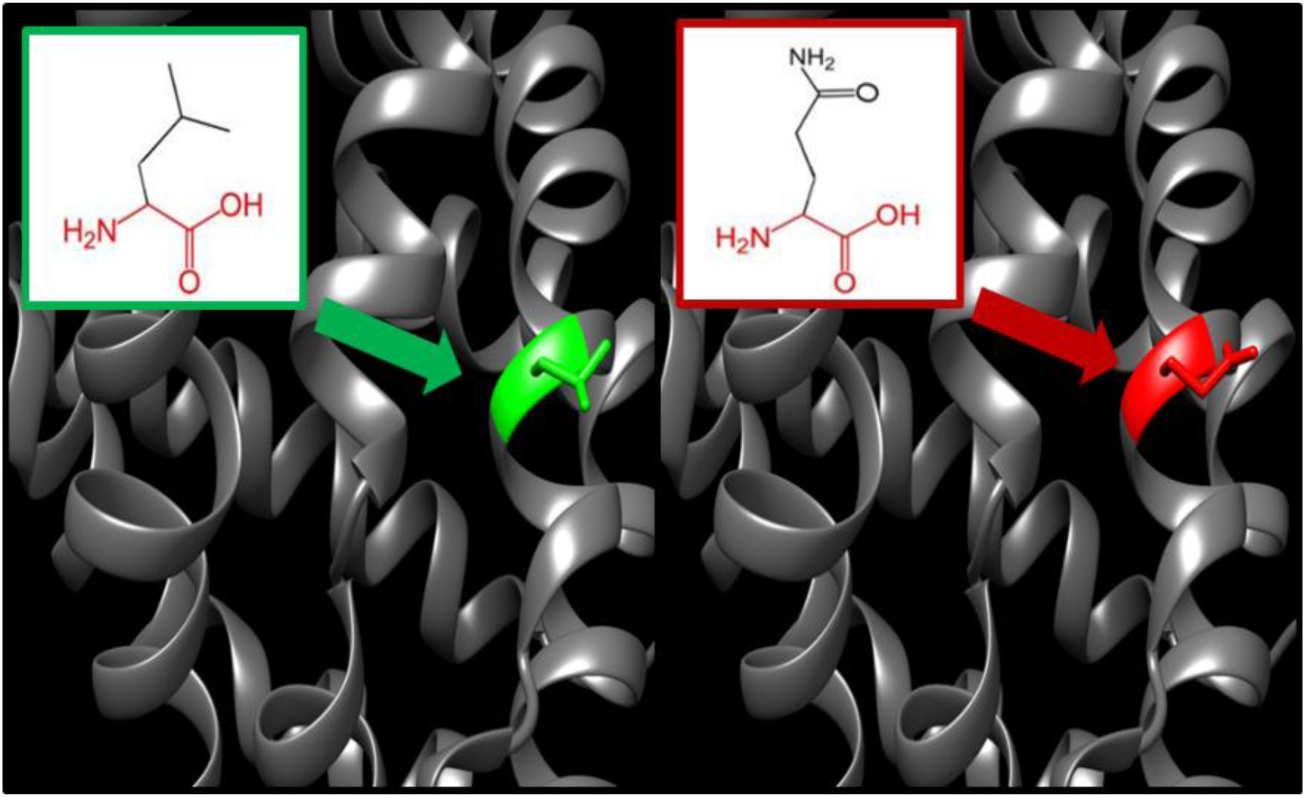
(L71Q) the amino acid Leucine (green) changes into a Glutamine (red) at position 71.

**Figure 7:**
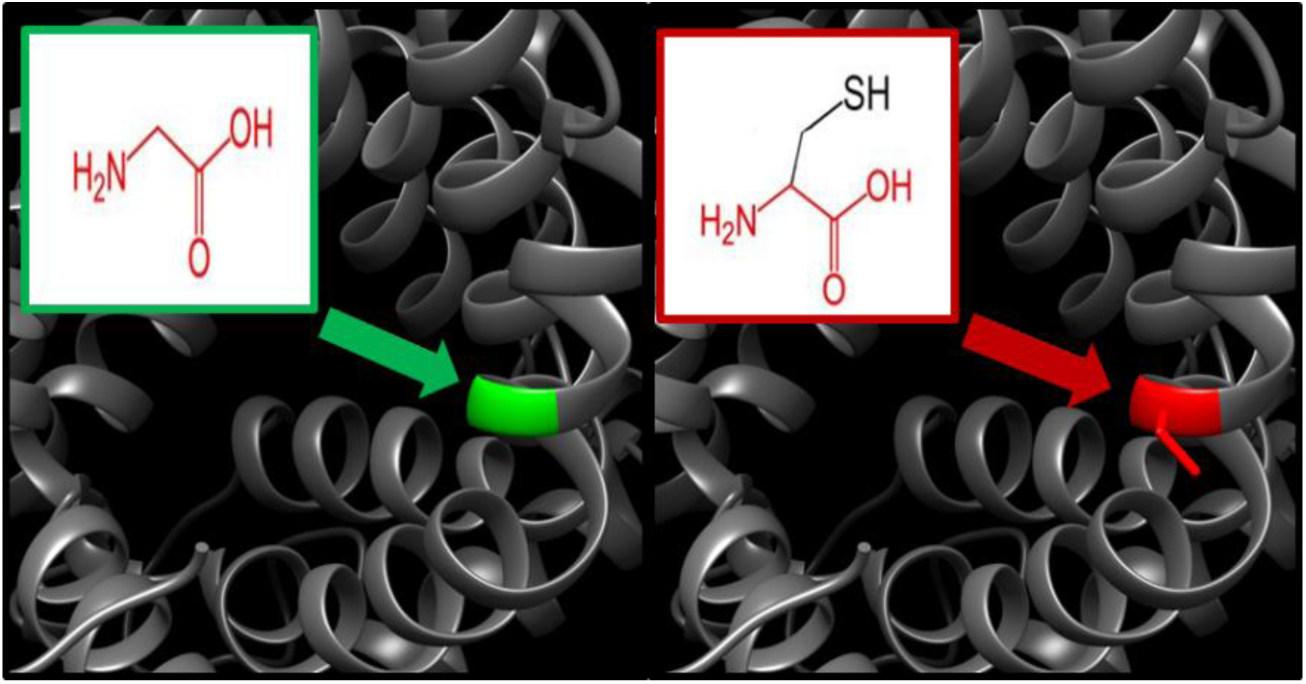
(G86C) the amino acid Glycine (green) changes into a Cysteine (red) at position 86.

**Figure 8:**
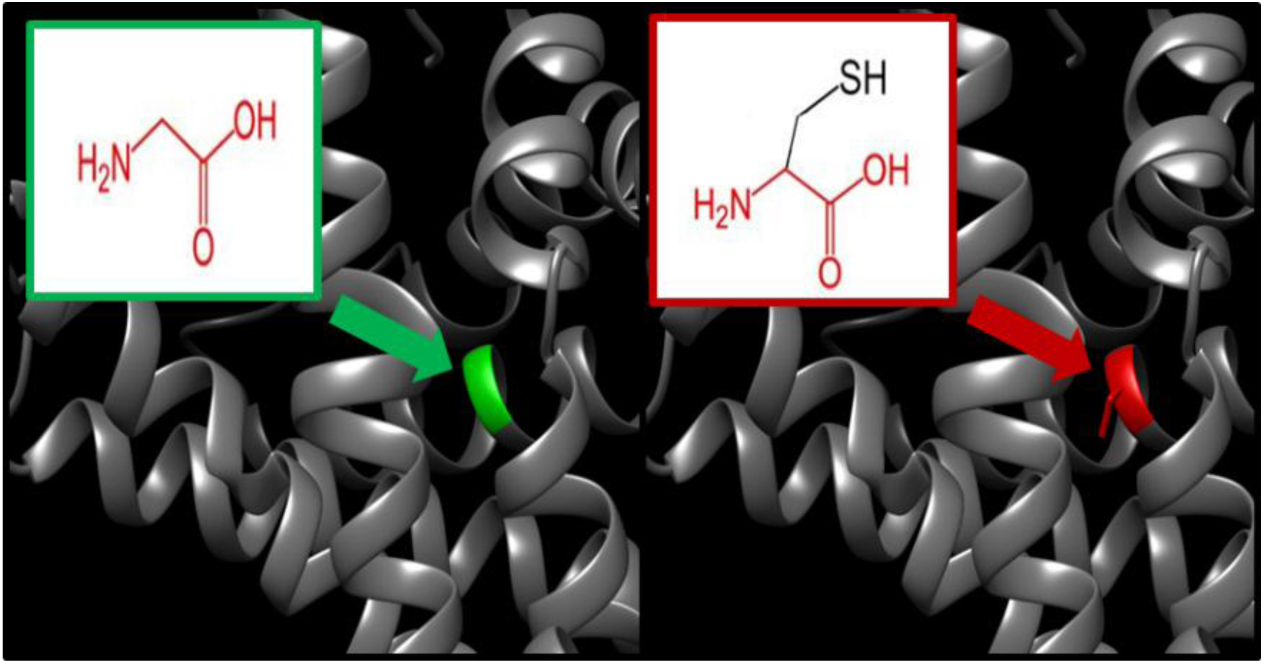
(G113C) the amino Acid Glycine (green) changes into a Cysteine (red) at position 113.

**Figure 9:**
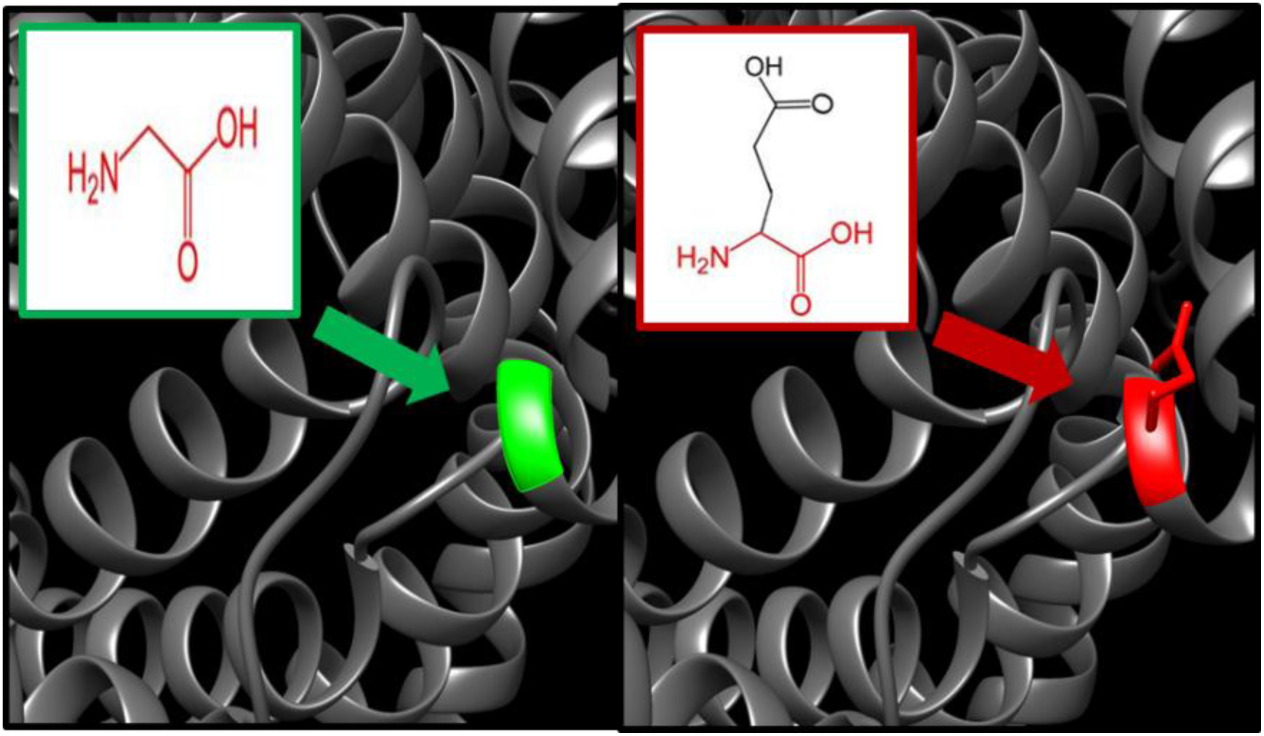
(G168E) the amino acid Glycine (green) changes into a Glutamic Acid (red) at position 168.

**Figure 10:**
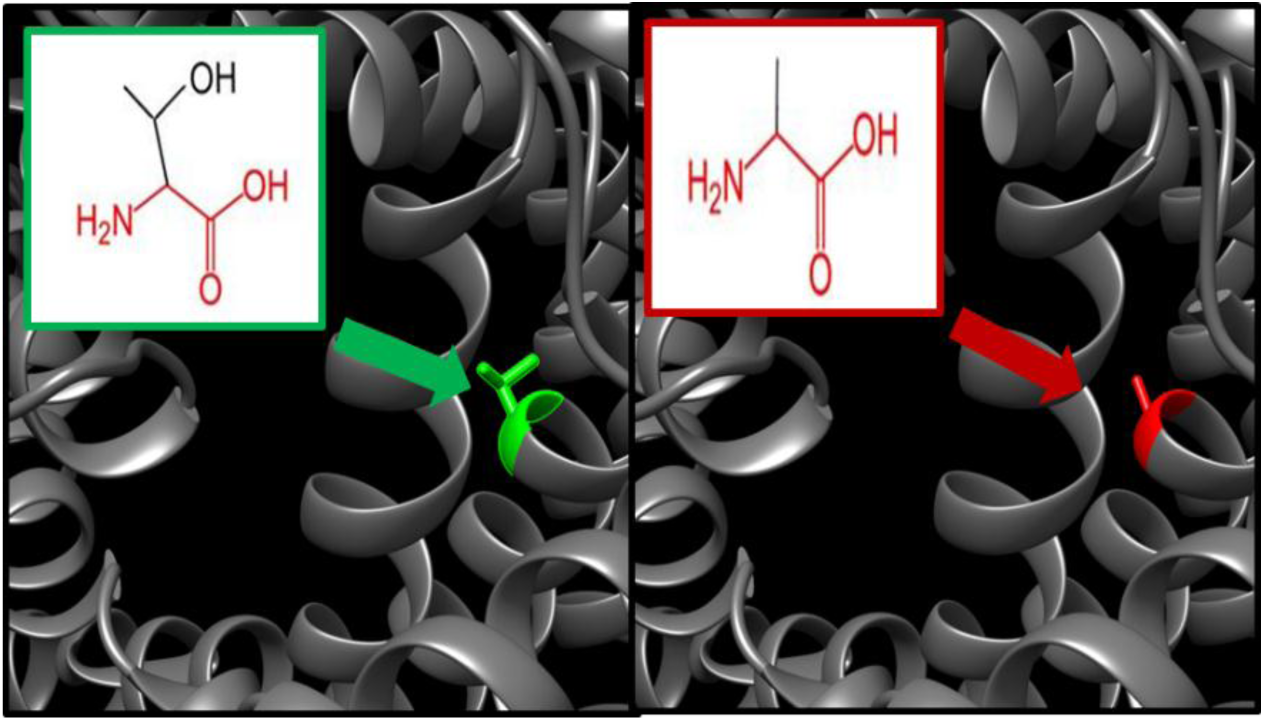
(T176A) the amino acid Threonine (green) changes into Alanine (red) at position 176.

**Figure 11:**
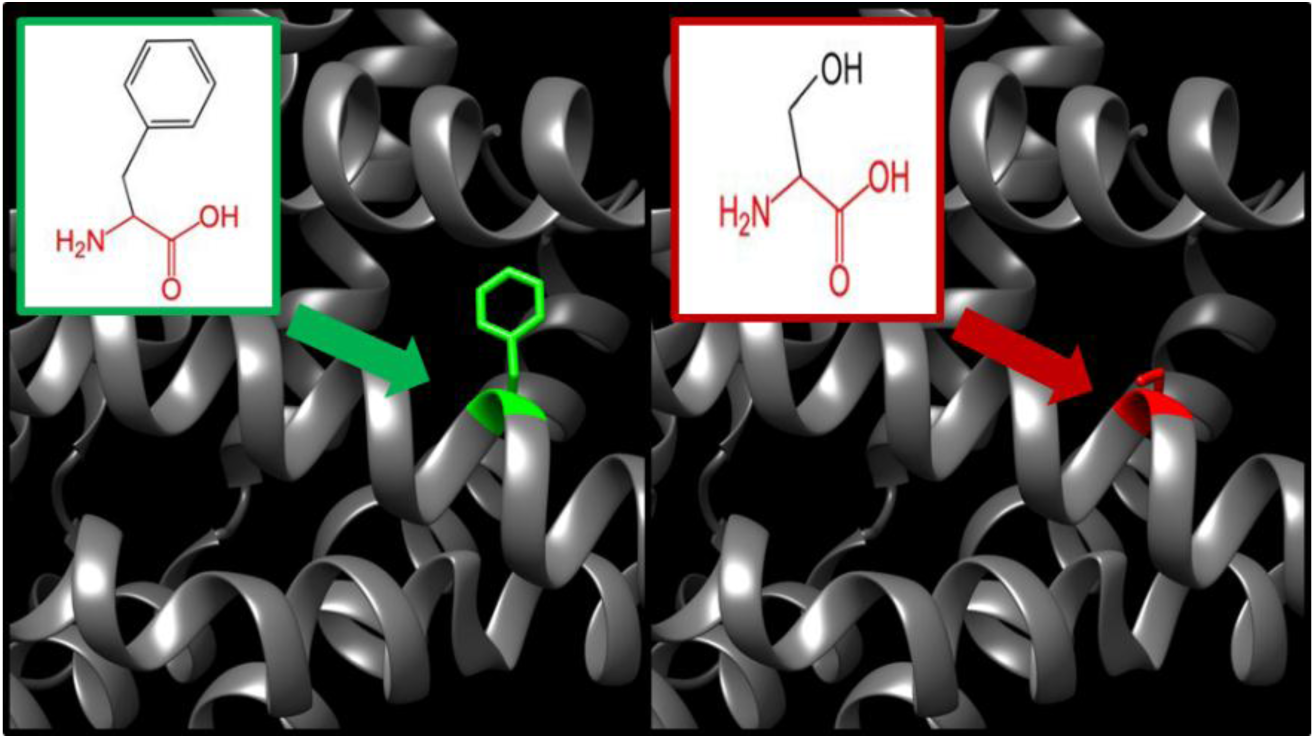
(F186S) the amino acid Phenylalanine (green) changes to Serine (red) at position 188.

**Figure 12:**
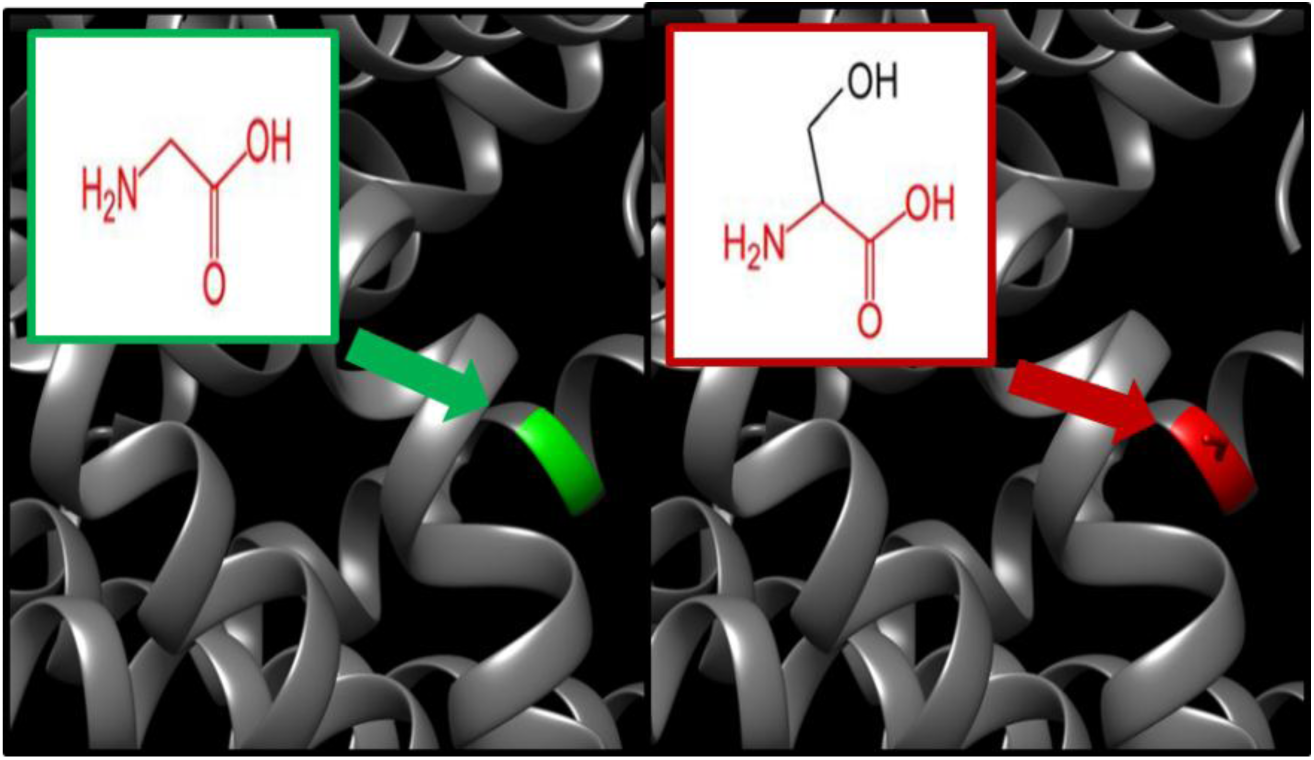
(G216S) the amino acid Glycine (green) changes to Serine 9 (red) at position 216.

**Figure 13:**
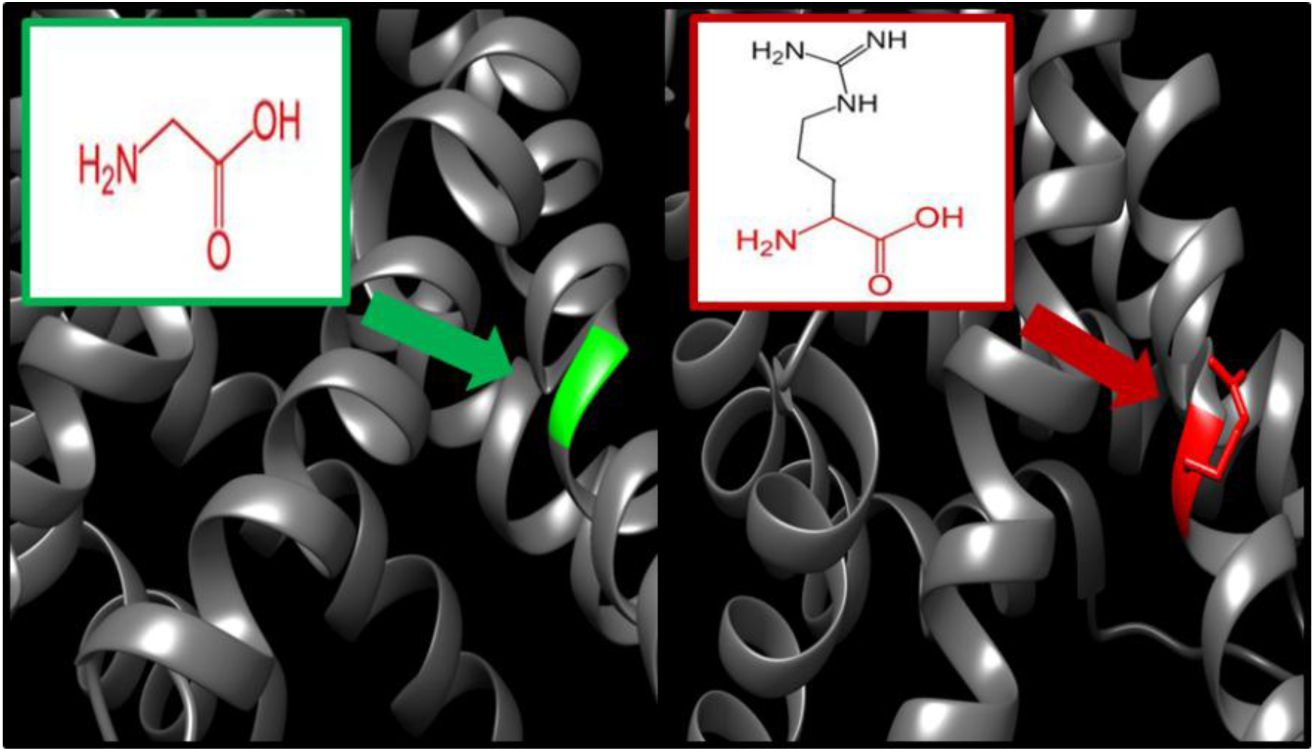
(G217R) the amino acid Glycine (green) change to Arginine (red) at position 217

**Figure 14:**
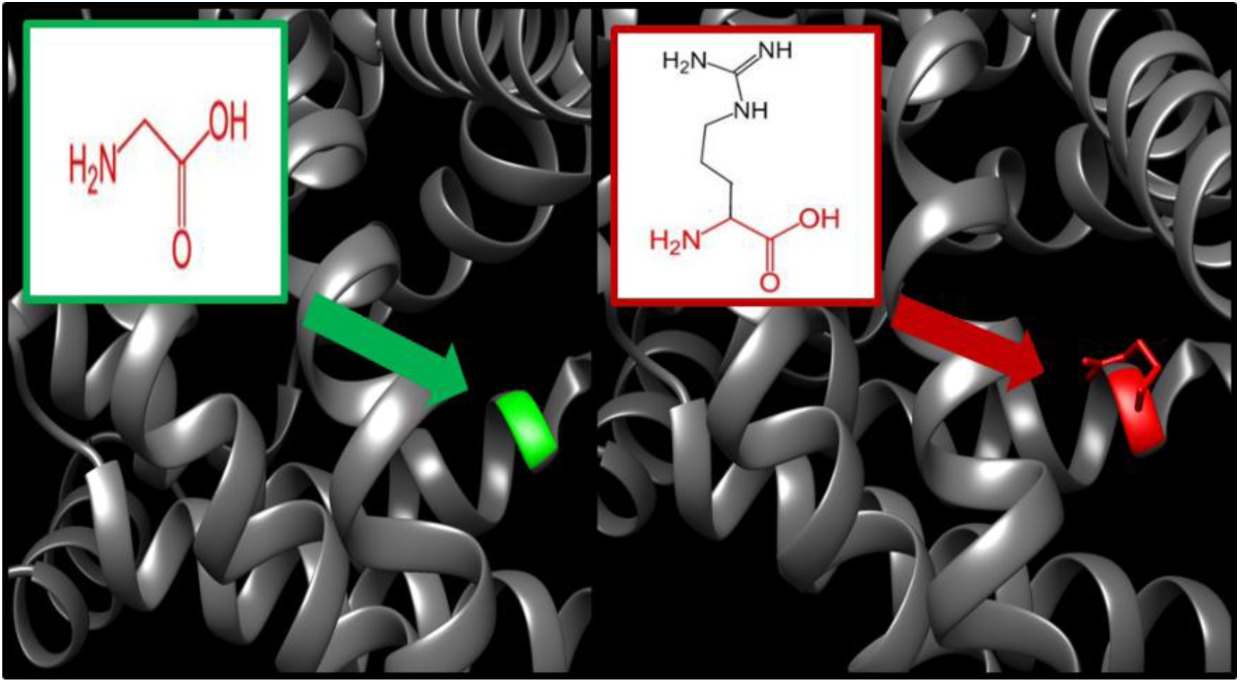
(G220R) the amino acid Glycine (green) change to Arginine (red) at position 220.

**Figure 15:**
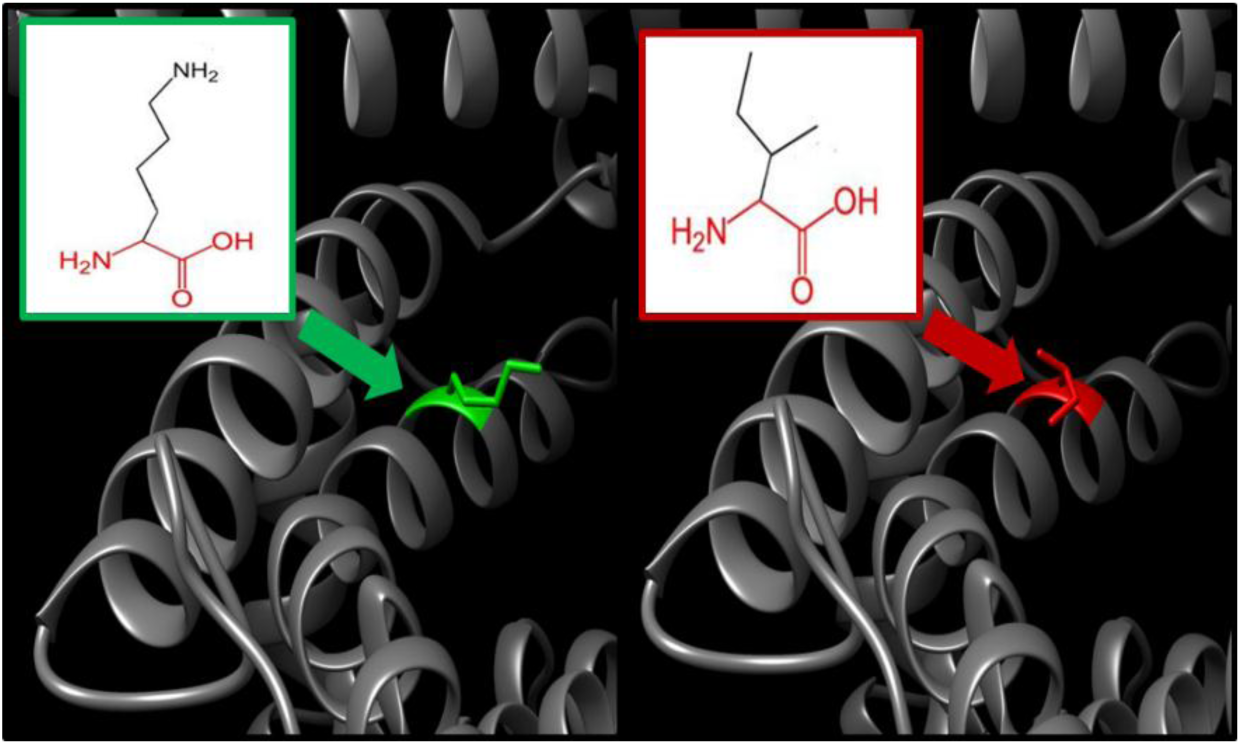
(K234I) the amino acid Lysine (green) changes to Isoleucine (red) at position 234.

**Figure 16:**
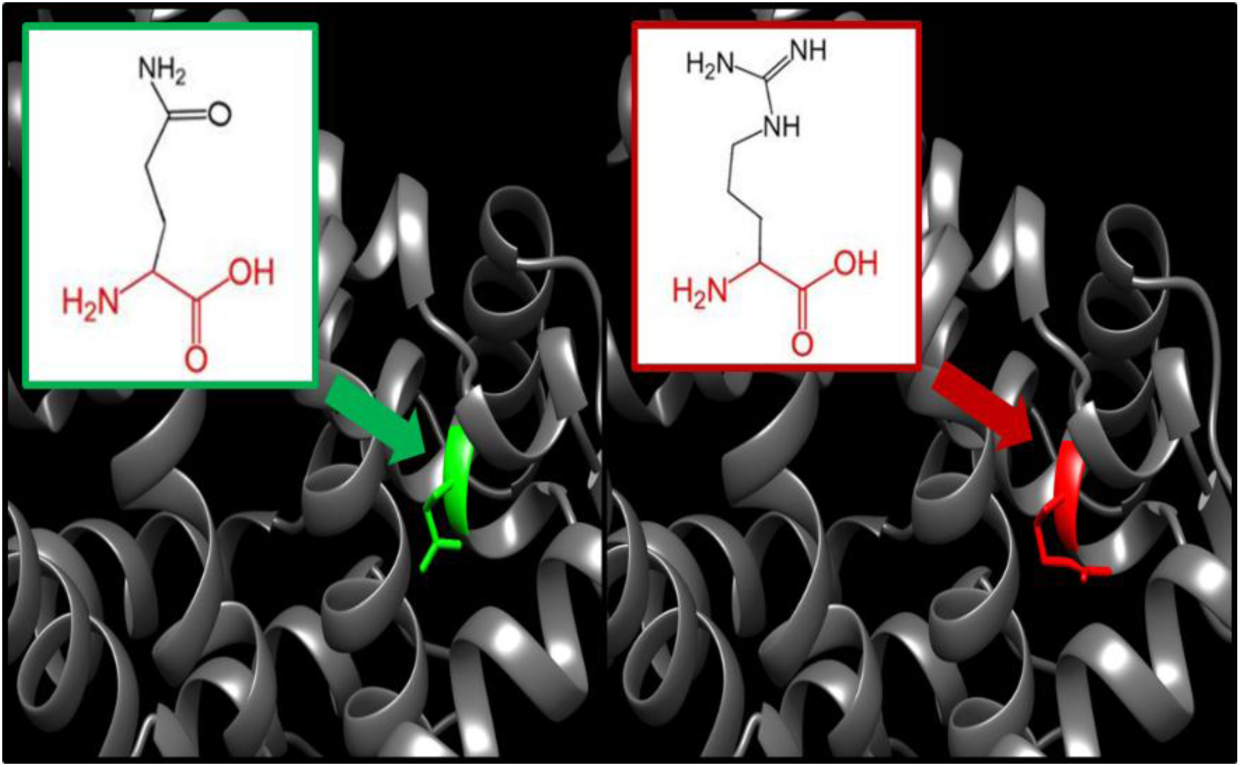
(Q238R) the amino acid Glutamine (green) changes to Arginine (red) at position 238.

**Figure 17:**
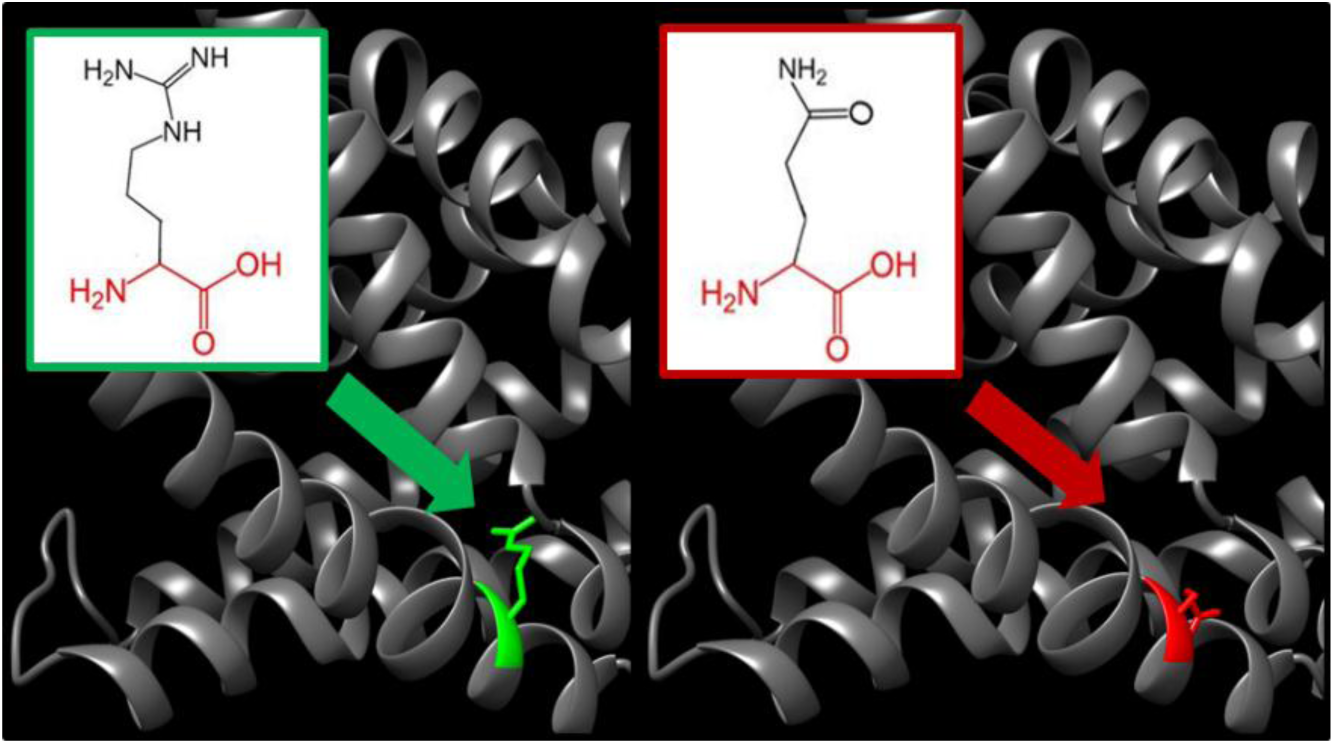
(R275Q) the amino acid Arginine (green) changes to Glutamine (red) at position 275.

**Figure 18:**
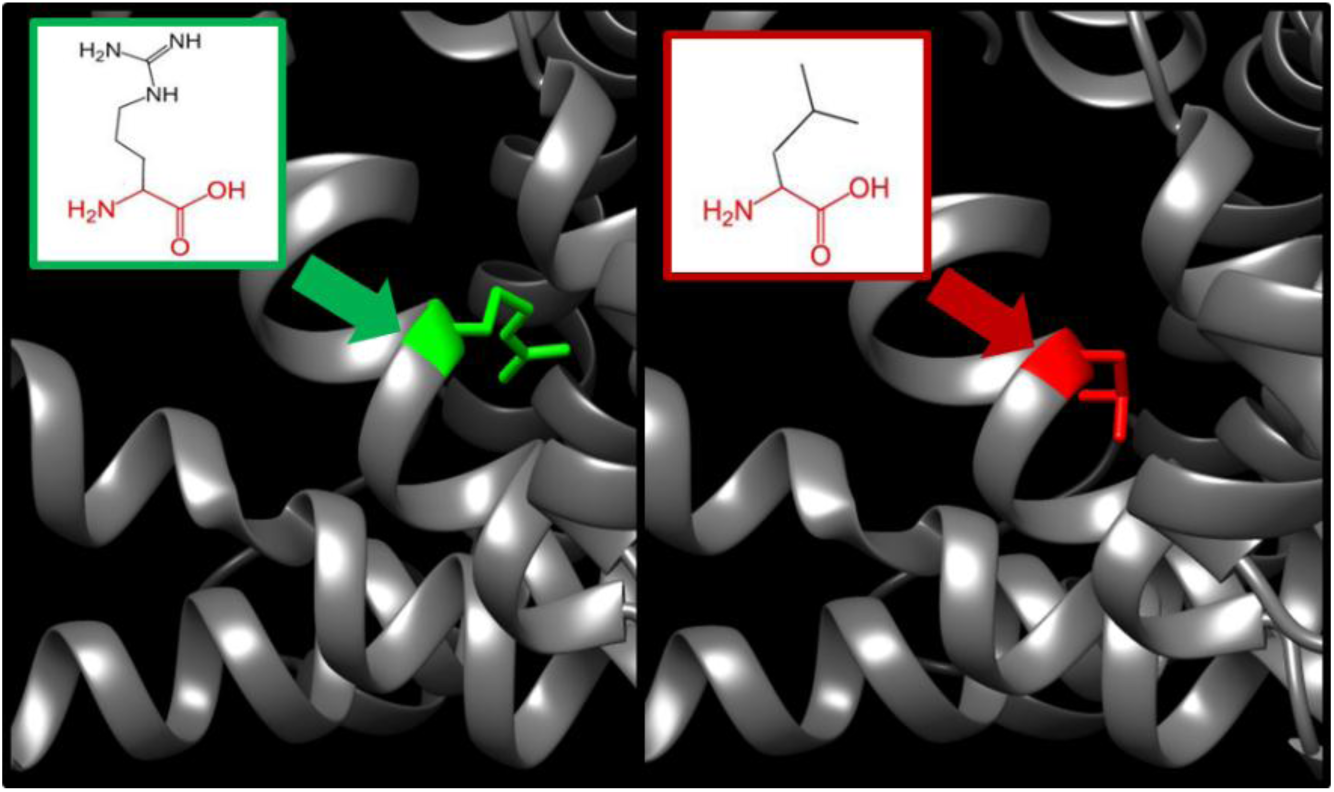
(R275L) the amino acid Arginine (green) change to Leucine (red) at position 275.

**Figure 19:**
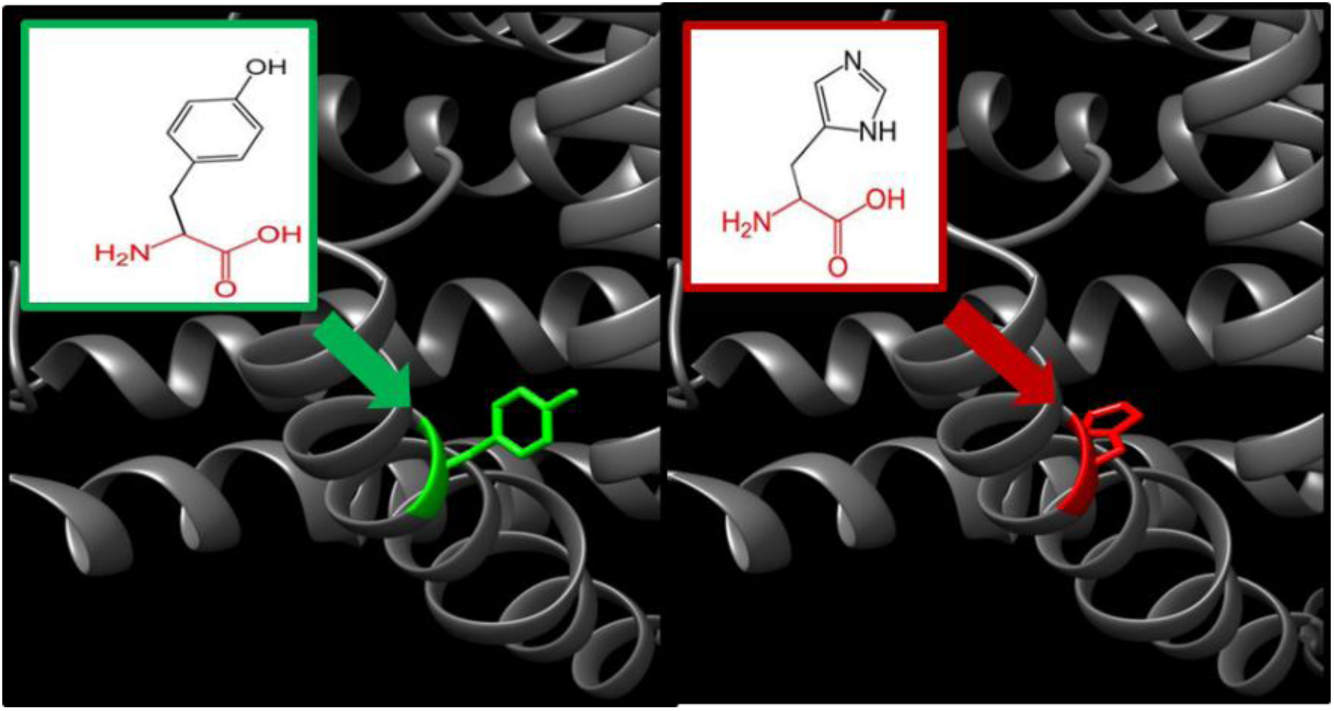
(Y287H) the amino acid Tyrosine (green) changes to Histidine (red) at position 287.

**Figure 20:**
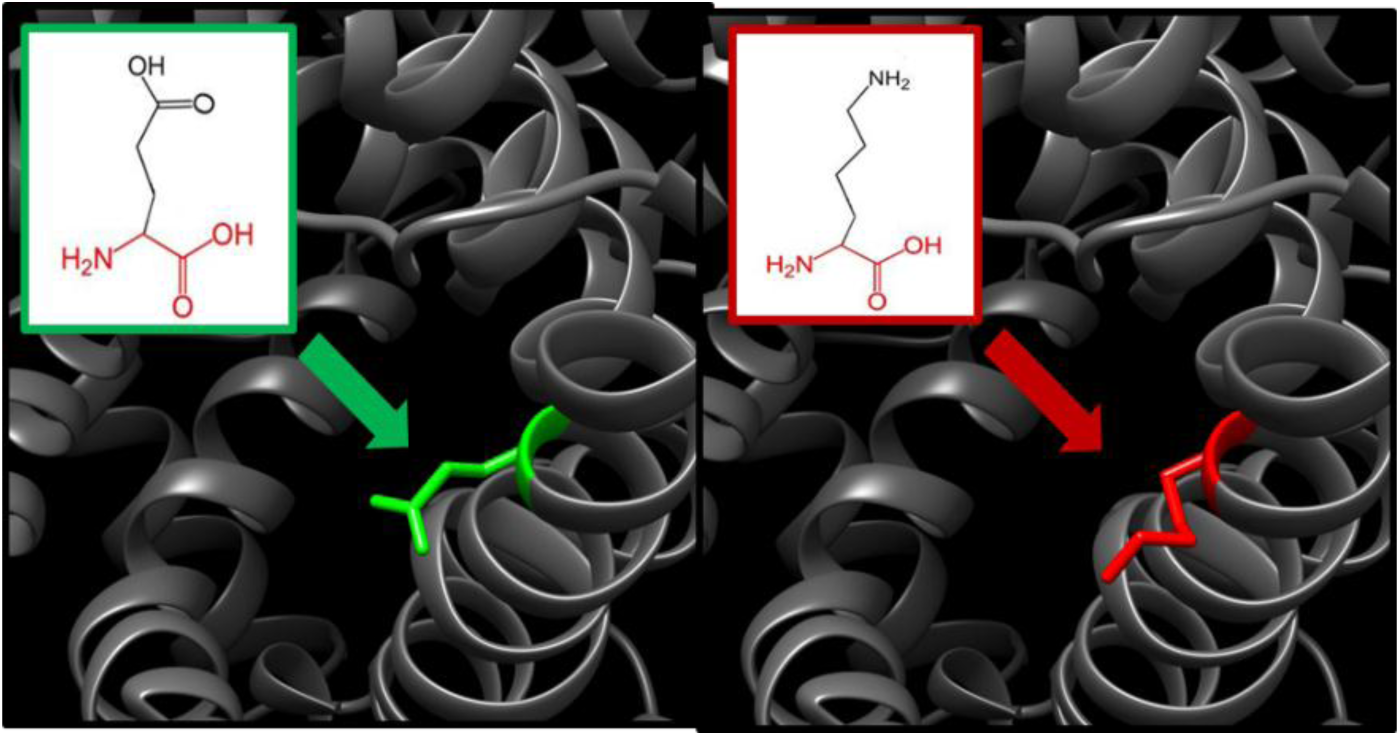
(E288K) the amino acid Glutamic Acid (green) changes to Lysine (red) at position 288.

**Figure 21:**
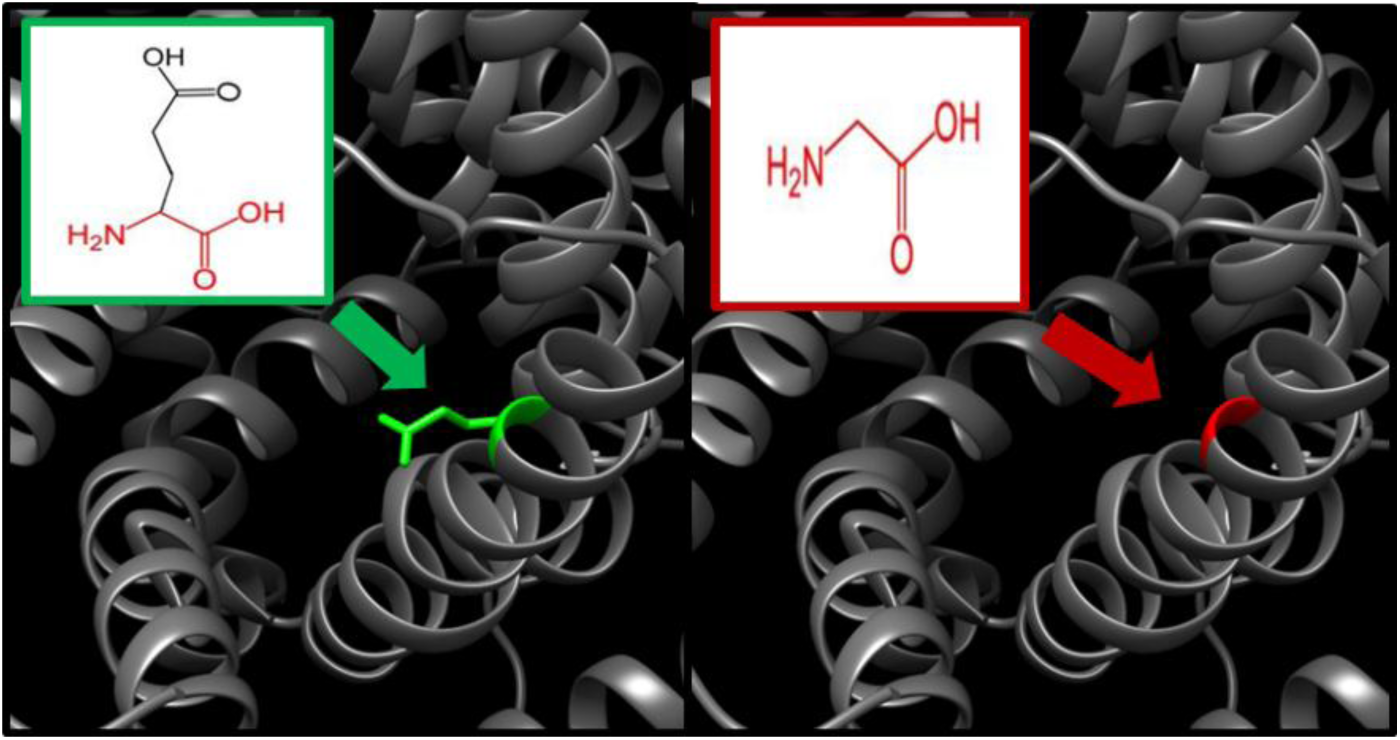
(E288G) the amino acid Glutamic Acid (green) changes into a Glycine (red) at position 288.

GeneMania disclosed that *SLC25A15* functions in the amide biosynthetic process, nitrogen cycle metabolic process, urea metabolic process, cellular amino acid catabolic process, inflammatory response, complement activation, cellular modified amino acid metabolic process, carboxylic acid catabolic process, organic acid catabolic process, protein activation cascade. Disruption of this function give rise to cytoplasmic ornithine aggregation, citrulline synthesis reduction, impaired ammonia detoxication and increased carbamoyl phosphate(1). The genes co-expressed with; share similar protein domain, or participate to achieve similar function were illustrated by GeneMANIA and shown in figure (22) Tables (5, and 6).

**Figure 22:**
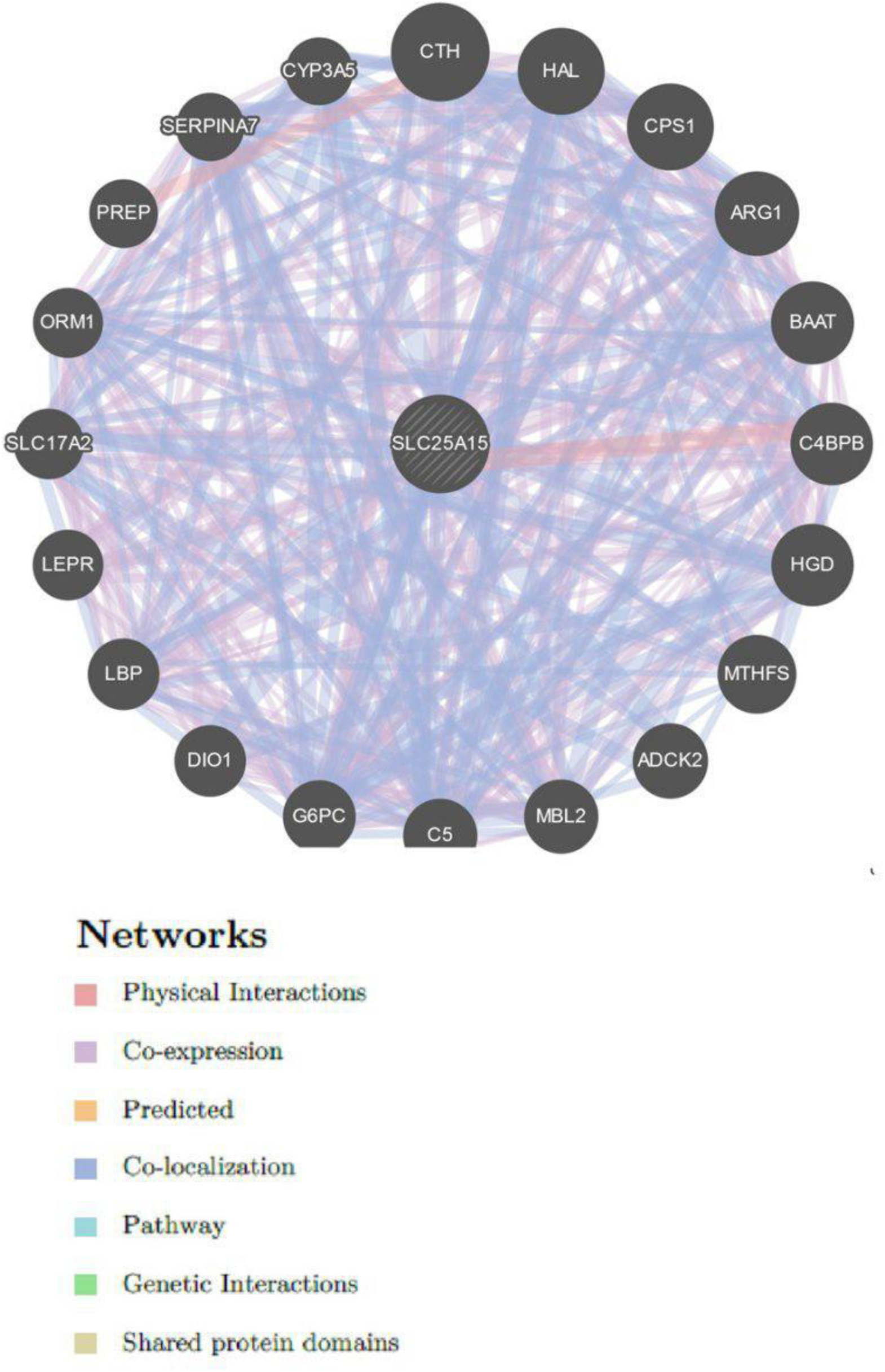
Interaction between *SLC25A15* and its related genes.

**Table (5):** *The SCL25A15* gene function and its appearance in network and genome:

| Function | FDR | Genes in network | Genes in genome |
| --- | --- | --- | --- |
| alpha-amino acid metabolic process | 1.90205E-06 | 7 | 142 |
| nitrogen cycle metabolic process | 0.001517377 | 3 | 12 |
| urea metabolic process | 0.001517377 | 3 | 11 |
| acute inflammatory response | 0.003303863 | 4 | 66 |
| cellular amino acid catabolic process | 0.009215468 | 4 | 90 |
| amide biosynthetic process | 0.022215719 | 3 | 35 |
| inflammatory response | 0.028672552 | 5 | 283 |
| complement activation | 0.031243953 | 3 | 43 |
| cellular modified amino acid metabolic process | 0.034067443 | 4 | 145 |
| carboxylic acid catabolic process | 0.042072255 | 4 | 161 |
| organic acid catabolic process | 0.042072255 | 4 | 161 |
| protein activation cascade | 0.059994966 | 3 | 61 |
| alpha-amino acid catabolic process | 0.067027879 | 3 | 65 |
| small molecule catabolic process | 0.075684161 | 4 | 199 |

**Table (6):** The gene co-expressed, shared domain and interaction with SCL25A15 gene network:

| Gene 1 | Gene 2 | Weight | Network group |
| --- | --- | --- | --- |
| *G6PC* | *ARG1* | 0.013130839 | Co-expression |
| *CTH* | *SLC25A15* | 0.026114333 | Co-expression |
| *ARG1* | *HAL* | 0.015584574 | Co-expression |
| *BAAT* | *HAL* | 0.010479293 | Co-expression |
| *BAAT* | *ARG1* | 0.010356814 | Co-expression |
| *C4BPB* | *BAAT* | 0.009120464 | Co-expression |
| *MTHFS* | *SLC25A15* | 0.025209978 | Co-expression |
| *MTHFS* | *CTH* | 0.023388306 | Co-expression |
| *MBL2* | *HAL* | 0.012773992 | Co-expression |
| *MBL2* | *HGD* | 0.00896276 | Co-expression |
| *C5* | *HAL* | 0.011430247 | Co-expression |
| *C5* | *ARG1* | 0.01082647 | Co-expression |
| *C5* | *BAAT* | 0.00760778 | Co-expression |
| *C5* | *C4BPB* | 0.009530854 | Co-expression |
| *C5* | *HGD* | 0.007940412 | Co-expression |
| *C5* | *MBL2* | 0.009273721 | Co-expression |
| *G6PC* | *HAL* | 0.011071949 | Co-expression |
| *G6PC* | *BAAT* | 0.007867279 | Co-expression |
| *G6PC* | *C4BPB* | 0.009524476 | Co-expression |
| *G6PC* | *HGD* | 0.007717506 | Co-expression |
| *G6PC* | *MBL2* | 0.009207115 | Co-expression |
| *G6PC* | *C5* | 0.008035515 | Co-expression |
| *LBP* | *HAL* | 0.009203292 | Co-expression |
| *LBP* | *ARG1* | 0.009164802 | Co-expression |
| *LBP* | *BAAT* | 0.00658665 | Co-expression |
| *LBP* | *HGD* | 0.006415015 | Co-expression |
| *LBP* | *MBL2* | 0.007615251 | Co-expression |
| *LBP* | *C5* | 0.006644955 | Co-expression |
| *LBP* | *G6PC* | 0.006901765 | Co-expression |
| *SLC17A2* | *HAL* | 0.010264788 | Co-expression |
| *SLC17A2* | *BAAT* | 0.007278958 | Co-expression |
| *SLC17A2* | *C4BPB* | 0.009022474 | Co-expression |
| *SLC17A2* | *HGD* | 0.007203176 | Co-expression |
| *SLC17A2* | *MBL2* | 0.008483202 | Co-expression |
| *SLC17A2* | *C5* | 0.007472467 | Co-expression |
| *SLC17A2* | *G6PC* | 0.007659466 | Co-expression |
| *SLC17A2* | *LBP* | 0.006347752 | Co-expression |
| *ORM1* | *HAL* | 0.009871504 | Co-expression |
| *ORM1* | *ARG1* | 0.009783008 | Co-expression |
| *ORM1* | *BAAT* | 0.007071007 | Co-expression |
| *ORM1* | *C4BPB* | 0.008490616 | Co-expression |
| *ORM1* | *MBL2* | 0.008121029 | Co-expression |
| *ORM1* | *C5* | 0.007148504 | Co-expression |
| *ORM1* | *G6PC* | 0.007398048 | Co-expression |
| *ORM1* | *LBP* | 0.006209759 | Co-expression |
| *ORM1* | *SLC17A2* | 0.006826424 | Co-expression |
| *PREP* | *SLC25A15* | 0.010442503 | Co-expression |
| *SERPINA7* | *MBL2* | 0.013247599 | Co-expression |
| *CPS1* | *HAL* | 0.01234286 | Co-expression |
| *ARG1* | *SLC25A15* | 0.005247516 | Co-expression |
| *ARG1* | *CTH* | 0.005402988 | Co-expression |
| *BAAT* | *SLC25A15* | 0.00938927 | Co-expression |
| *BAAT* | *HAL* | 0.008351558 | Co-expression |
| *BAAT* | *ARG1* | 0.004348387 | Co-expression |
| *C4BPB* | *SLC25A15* | 0.009231251 | Co-expression |
| *C4BPB* | *HAL* | 0.00814306 | Co-expression |
| *C4BPB* | *BAAT* | 0.007613069 | Co-expression |
| *HGD* | *HAL* | 0.013029266 | Co-expression |
| *HGD* | *ARG1* | 0.006771524 | Co-expression |
| *HGD* | *BAAT* | 0.012214398 | Co-expression |
| *HGD* | *C4BPB* | 0.011807794 | Co-expression |
| *MBL2* | *SLC25A15* | 0.008736178 | Co-expression |
| *MBL2* | *HAL* | 0.007751754 | Co-expression |
| *MBL2* | *BAAT* | 0.007231998 | Co-expression |
| *MBL2* | *C4BPB* | 0.00703362 | Co-expression |
| *MBL2* | *HGD* | 0.011195325 | Co-expression |
| *C5* | *SLC25A15* | 0.006322735 | Co-expression |
| *C5* | *HAL* | 0.005552603 | Co-expression |
| *C5* | *ARG1* | 0.002911721 | Co-expression |
| *C5* | *BAAT* | 0.005179715 | Co-expression |
| *C5* | *C4BPB* | 0.005058356 | Co-expression |
| *C5* | *HGD* | 0.008068559 | Co-expression |
| *C5* | *MBL2* | 0.004826631 | Co-expression |
| *G6PC* | *CTH* | 0.006566833 | Co-expression |
| *G6PC* | *CPS1* | 0.007564845 | Co-expression |
| *G6PC* | *ARG1* | 0.002987566 | Co-expression |
| *DIO1* | *ARG1* | 0.003676503 | Co-expression |
| *DIO1* | *C5* | 0.004387379 | Co-expression |
| *DIO1* | *G6PC* | 0.004473042 | Co-expression |
| *LBP* | *BAAT* | 0.009965218 | Co-expression |
| *LBP* | *C4BPB* | 0.009667832 | Co-expression |
| *LBP* | *MBL2* | 0.009191572 | Co-expression |
| *LBP* | *C5* | 0.006629785 | Co-expression |
| *SLC17A2* | *SLC25A15* | 0.007915142 | Co-expression |
| *SLC17A2* | *HAL* | 0.007040063 | Co-expression |
| *SLC17A2* | *CPS1* | 0.009285137 | Co-expression |
| *SLC17A2* | *BAAT* | 0.006526508 | Co-expression |
| *SLC17A2* | *C4BPB* | 0.006351683 | Co-expression |
| *SLC17A2* | *HGD* | 0.010178178 | Co-expression |
| *SLC17A2* | *MBL2* | 0.006092981 | Co-expression |
| *SLC17A2* | *C5* | 0.004395986 | Co-expression |
| *SLC17A2* | *DIO1* | 0.005542256 | Co-expression |
| *SLC17A2* | *LBP* | 0.008360203 | Co-expression |
| *ORM1* | *SLC25A15* | 0.009182534 | Co-expression |
| *ORM1* | *C5* | 0.005041816 | Co-expression |
| *ORM1* | *G6PC* | 0.005162204 | Co-expression |
| *SERPINA7* | *SLC25A15* | 0.007883273 | Co-expression |
| *SERPINA7* | *HAL* | 0.006906412 | Co-expression |
| *SERPINA7* | *CPS1* | 0.009286882 | Co-expression |
| *SERPINA7* | *ARG1* | 0.003641196 | Co-expression |
| *SERPINA7* | *C4BPB* | 0.006316919 | Co-expression |
| *SERPINA7* | *MBL2* | 0.005971783 | Co-expression |
| *SERPINA7* | *C5* | 0.00436016 | Co-expression |
| *SERPINA7* | *DIO1* | 0.005475692 | Co-expression |
| *SERPINA7* | *SLC17A2* | 0.005462843 | Co-expression |
| *SERPINA7* | *ORM1* | 0.006366141 | Co-expression |
| *CYP3A5* | *C4BPB* | 0.009888263 | Co-expression |
| *CYP3A5* | *MBL2* | 0.009454532 | Co-expression |
| *CYP3A5* | *C5* | 0.006786631 | Co-expression |
| *CYP3A5* | *SLC17A2* | 0.008575552 | Co-expression |
| *CYP3A5* | *ORM1* | 0.009884344 | Co-expression |
| *CYP3A5* | *SERPINA7* | 0.008449458 | Co-expression |
| *CTH* | *SLC25A15* | 0.017681984 | Co-expression |
| *HAL* | *SLC25A15* | 0.011282469 | Co-expression |
| *HAL* | *CTH* | 0.018665954 | Co-expression |
| *CPS1* | *SLC25A15* | 0.014783018 | Co-expression |
| *CPS1* | *CTH* | 0.021131555 | Co-expression |
| *CPS1* | *HAL* | 0.01605092 | Co-expression |
| *ARG1* | *SLC25A15* | 0.01022149 | Co-expression |
| *ARG1* | *CTH* | 0.014448987 | Co-expression |
| *ARG1* | *CPS1* | 0.014791401 | Co-expression |
| *BAAT* | *SLC25A15* | 0.007653142 | Co-expression |
| *BAAT* | *CPS1* | 0.011089044 | Co-expression |
| *BAAT* | *ARG1* | 0.009111863 | Co-expression |
| *C4BPB* | *CTH* | 0.007979206 | Co-expression |
| *HGD* | *SLC25A15* | 0.009968358 | Co-expression |
| *HGD* | *CTH* | 0.011085113 | Co-expression |
| *HGD* | *HAL* | 0.008449321 | Co-expression |
| *HGD* | *CPS1* | 0.012882358 | Co-expression |
| *HGD* | *ARG1* | 0.009243058 | Co-expression |
| *HGD* | *BAAT* | 0.007479178 | Co-expression |
| *MTHFS* | *C4BPB* | 0.009367498 | Co-expression |
| *C5* | *HAL* | 0.007322867 | Co-expression |
| *C5* | *MBL2* | 0.010361436 | Co-expression |
| *G6PC* | *SLC25A15* | 0.016148888 | Co-expression |
| *G6PC* | *CTH* | 0.022236109 | Co-expression |
| *G6PC* | *HAL* | 0.01545969 | Co-expression |
| *G6PC* | *CPS1* | 0.01768735 | Co-expression |
| *G6PC* | *ARG1* | 0.012720793 | Co-expression |
| *G6PC* | *BAAT* | 0.009469871 | Co-expression |
| *G6PC* | *HGD* | 0.011001362 | Co-expression |
| *LBP* | *C4BPB* | 0.003940297 | Co-expression |
| *LEPR* | *SLC25A15* | 0.015385044 | Co-expression |
| *LEPR* | *HAL* | 0.018157363 | Co-expression |
| *LEPR* | *CPS1* | 0.021792563 | Co-expression |
| *LEPR* | *ARG1* | 0.012319317 | Co-expression |
| *LEPR* | *HGD* | 0.012408955 | Co-expression |
| *LEPR* | *G6PC* | 0.026779622 | Co-expression |
| *SLC17A2* | *C4BPB* | 0.006601628 | Co-expression |
| *SLC17A2* | *MTHFS* | 0.011673527 | Co-expression |
| *SLC17A2* | *LBP* | 0.004327369 | Co-expression |
| *ORM1* | *C4BPB* | 0.008727606 | Co-expression |
| *ORM1* | *C5* | 0.007064989 | Co-expression |
| *PREP* | *SLC25A15* | 0.013786578 | Co-expression |
| *SERPINA7* | *C5* | 0.005918165 | Co-expression |
| *SERPINA7* | *ORM1* | 0.010403195 | Co-expression |
| *HGD* | *CPS1* | 0.020025367 | Co-expression |
| *DIO1* | *MBL2* | 0.014330983 | Co-expression |
| *LBP* | *G6PC* | 0.0091696 | Co-expression |
| *CPS1* | *SLC25A15* | 0.010167049 | Co-expression |
| *CTH* | *SLC25A15* | 0.022405656 | Co-expression |
| *HAL* | *SLC25A15* | 0.024649095 | Co-expression |
| *CPS1* | *SLC25A15* | 0.011441315 | Co-expression |
| *CPS1* | *CTH* | 0.011585185 | Co-expression |
| *CPS1* | *HAL* | 0.012319576 | Co-expression |
| *ARG1* | *SLC25A15* | 0.020499837 | Co-expression |
| *BAAT* | *SLC25A15* | 0.01333323 | Co-expression |
| *BAAT* | *CTH* | 0.01358016 | Co-expression |
| *BAAT* | *CPS1* | 0.007241853 | Co-expression |
| *BAAT* | *ARG1* | 0.011780513 | Co-expression |
| *C4BPB* | *SLC25A15* | 0.010655208 | Co-expression |
| *C4BPB* | *CTH* | 0.010559047 | Co-expression |
| *C4BPB* | *HAL* | 0.011841011 | Co-expression |
| *C4BPB* | *CPS1* | 0.006141116 | Co-expression |
| *HGD* | *CPS1* | 0.009511196 | Co-expression |
| *HGD* | *BAAT* | 0.011432248 | Co-expression |
| *MBL2* | *SLC25A15* | 0.017125212 | Co-expression |
| *MBL2* | *CTH* | 0.016398663 | Co-expression |
| *MBL2* | *HAL* | 0.018915584 | Co-expression |
| *MBL2* | *CPS1* | 0.009279709 | Co-expression |
| *MBL2* | *C4BPB* | 0.008994715 | Co-expression |
| *C5* | *SLC25A15* | 0.008921804 | Co-expression |
| *C5* | *CTH* | 0.008959468 | Co-expression |
| *C5* | *HAL* | 0.009587029 | Co-expression |
| *C5* | *CPS1* | 0.005092065 | Co-expression |
| *C5* | *BAAT* | 0.005769644 | Co-expression |
| *C5* | *C4BPB* | 0.004815416 | Co-expression |
| *C5* | *HGD* | 0.007569547 | Co-expression |
| *C5* | *MBL2* | 0.007248661 | Co-expression |
| *G6PC* | *CTH* | 0.021676406 | Co-expression |
| *G6PC* | *CPS1* | 0.010997106 | Co-expression |
| *G6PC* | *BAAT* | 0.012542194 | Co-expression |
| *G6PC* | *C4BPB* | 0.009974403 | Co-expression |
| *G6PC* | *HGD* | 0.017824262 | Co-expression |
| *G6PC* | *C5* | 0.00849682 | Co-expression |
| *LBP* | *SLC25A15* | 0.018759409 | Co-expression |
| *LBP* | *HAL* | 0.02087546 | Co-expression |
| *LBP* | *CPS1* | 0.009439292 | Co-expression |
| *LBP* | *ARG1* | 0.018111857 | Co-expression |
| *LBP* | *BAAT* | 0.010925728 | Co-expression |
| *LBP* | *C4BPB* | 0.009051763 | Co-expression |
| *LBP* | *MBL2* | 0.014943773 | Co-expression |
| *LBP* | *C5* | 0.007415157 | Co-expression |
| *LEPR* | *SLC25A15* | 0.025149433 | Co-expression |
| *LEPR* | *CTH* | 0.02639034 | Co-expression |
| *LEPR* | *CPS1* | 0.012923609 | Co-expression |
| *LEPR* | *BAAT* | 0.015909543 | Co-expression |
| *LEPR* | *C5* | 0.010308038 | Co-expression |
| *LEPR* | *LBP* | 0.020950127 | Co-expression |
| *SLC17A2* | *C4BPB* | 0.005922639 | Co-expression |
| *SLC17A2* | *C5* | 0.00483539 | Co-expression |
| *ORM1* | *SLC25A15* | 0.012226423 | Co-expression |
| *ORM1* | *CTH* | 0.012432948 | Co-expression |
| *ORM1* | *HAL* | 0.013200064 | Co-expression |
| *ORM1* | *CPS1* | 0.006929017 | Co-expression |
| *ORM1* | *BAAT* | 0.007673539 | Co-expression |
| *ORM1* | *C4BPB* | 0.00653899 | Co-expression |
| *ORM1* | *HGD* | 0.010120259 | Co-expression |
| *ORM1* | *MBL2* | 0.009900892 | Co-expression |
| *ORM1* | *C5* | 0.005402325 | Co-expression |
| *ORM1* | *G6PC* | 0.011815394 | Co-expression |
| *ORM1* | *LBP* | 0.010073603 | Co-expression |
| *ORM1* | *LEPR* | 0.013810683 | Co-expression |
| *SERPINA7* | *SLC25A15* | 0.012462362 | Co-expression |
| *SERPINA7* | *HAL* | 0.01432913 | Co-expression |
| *SERPINA7* | *CPS1* | 0.00686181 | Co-expression |
| *SERPINA7* | *ARG1* | 0.01146561 | Co-expression |
| *SERPINA7* | *C4BPB* | 0.00677565 | Co-expression |
| *SERPINA7* | *MBL2* | 0.010809767 | Co-expression |
| *SERPINA7* | *C5* | 0.005448063 | Co-expression |
| *SERPINA7* | *LBP* | 0.011217129 | Co-expression |
| *SERPINA7* | *SLC17A2* | 0.006874819 | Co-expression |
| *CYP3A5* | *CPS1* | 0.01207218 | Co-expression |
| *CYP3A5* | *C4BPB* | 0.011685826 | Co-expression |
| *CYP3A5* | *C5* | 0.009578024 | Co-expression |
| *CYP3A5* | *SLC17A2* | 0.011991991 | Co-expression |
| *CYP3A5* | *ORM1* | 0.012806383 | Co-expression |
| *SLC17A2* | *G6PC* | 0.014184606 | Co-expression |
| *SERPINA7* | *C5* | 0.019346774 | Co-expression |
| *CYP3A5* | *SERPINA7* | 0.010969018 | Co-expression |
| *HGD* | *SLC25A15* | 0.006551247 | Co-expression |
| *PREP* | *SLC25A15* | 0.012394655 | Co-expression |
| *SERPINA7* | *G6PC* | 0.017327776 | Co-expression |
| *CPS1* | *HAL* | 0.00694992 | Co-expression |
| *ARG1* | *CTH* | 0.006446471 | Co-expression |
| *ARG1* | *HAL* | 0.010185538 | Co-expression |
| *ARG1* | *CPS1* | 0.006081839 | Co-expression |
| *BAAT* | *ARG1* | 0.01191919 | Co-expression |
| *C4BPB* | *CPS1* | 0.00363932 | Co-expression |
| *HGD* | *CPS1* | 0.008691518 | Co-expression |
| *HGD* | *ARG1* | 0.010913373 | Co-expression |
| *MTHFS* | *HAL* | 0.007126623 | Co-expression |
| *C5* | *C4BPB* | 0.004553121 | Co-expression |
| *LBP* | *ARG1* | 0.006003315 | Co-expression |
| *LBP* | *C4BPB* | 0.00376871 | Co-expression |
| *LBP* | *C5* | 0.005731973 | Co-expression |
| *LEPR* | *CPS1* | 0.010181712 | Co-expression |
| *LEPR* | *ARG1* | 0.012883171 | Co-expression |
| *LEPR* | *HGD* | 0.019583575 | Co-expression |
| *SERPINA7* | *C4BPB* | 0.006879414 | Co-expression |
| *SERPINA7* | *C5* | 0.009770601 | Co-expression |
| *SERPINA7* | *DIO1* | 0.018938221 | Co-expression |
| *SERPINA7* | *SLC17A2* | 0.021433447 | Co-expression |
| *DIO1* | *C4BPB* | 0.01167786 | Co-expression |
| *DIO1* | *CTH* | 0.004786647 | Co-expression |
| *DIO1* | *MBL2* | 0.007709226 | Co-expression |
| *DIO1* | *C5* | 0.006562013 | Co-expression |
| *HAL* | *CTH* | 0.00904644 | Co-localization |
| *ARG1* | *CTH* | 0.007396141 | Co-localization |
| *ARG1* | *HAL* | 0.011341748 | Co-localization |
| *C4BPB* | *CTH* | 0.00882447 | Co-localization |
| *C4BPB* | *HAL* | 0.009825678 | Co-localization |
| *MBL2* | *ARG1* | 0.011475822 | Co-localization |
| *C5* | *CTH* | 0.009882752 | Co-localization |
| *C5* | *HAL* | 0.012378897 | Co-localization |
| *C5* | *ARG1* | 0.011783864 | Co-localization |
| *C5* | *MBL2* | 0.015168445 | Co-localization |
| *G6PC* | *CTH* | 0.006479372 | Co-localization |
| *G6PC* | *HAL* | 0.007433198 | Co-localization |
| *G6PC* | *ARG1* | 0.006618402 | Co-localization |
| *G6PC* | *C4BPB* | 0.008764179 | Co-localization |
| *G6PC* | *HGD* | 0.011694877 | Co-localization |
| *SERPINA7* | *CTH* | 0.007116361 | Co-localization |
| *SERPINA7* | *HAL* | 0.008624428 | Co-localization |
| *SERPINA7* | *ARG1* | 0.007838565 | Co-localization |
| *SERPINA7* | *C4BPB* | 0.009594075 | Co-localization |
| *SERPINA7* | *MBL2* | 0.011912366 | Co-localization |
| *SERPINA7* | *C5* | 0.012075914 | Co-localization |
| *SERPINA7* | *G6PC* | 0.007159275 | Co-localization |
| *HAL* | *SLC25A15* | 0.014272561 | Co-localization |
| *CPS1* | *SLC25A15* | 0.013714614 | Co-localization |
| *ARG1* | *SLC25A15* | 0.01310207 | Co-localization |
| *ARG1* | *CPS1* | 0.00861983 | Co-localization |
| *BAAT* | *SLC25A15* | 0.013363565 | Co-localization |
| *BAAT* | *CPS1* | 0.008450216 | Co-localization |
| *BAAT* | *ARG1* | 0.008598406 | Co-localization |
| *HGD* | *SLC25A15* | 0.022047134 | Co-localization |
| *HGD* | *HAL* | 0.010812771 | Co-localization |
| *MTHFS* | *SLC25A15* | 0.008358937 | Co-localization |
| *MTHFS* | *HAL* | 0.004322442 | Co-localization |
| *MTHFS* | *C4BPB* | 0.009622298 | Co-localization |
| *MTHFS* | *HGD* | 0.007224194 | Co-localization |
| *ADCK2* | *SLC25A15* | 0.026127955 | Co-localization |
| *ADCK2* | *BAAT* | 0.010211586 | Co-localization |
| *ADCK2* | *MTHFS* | 0.00685143 | Co-localization |
| *MBL2* | *SLC25A15* | 0.011515018 | Co-localization |
| *MBL2* | *C4BPB* | 0.012219642 | Co-localization |
| *MBL2* | *MTHFS* | 0.003996356 | Co-localization |
| *MBL2* | *ADCK2* | 0.009659463 | Co-localization |
| *C5* | *SLC25A15* | 0.01863199 | Co-localization |
| *C5* | *HAL* | 0.009531116 | Co-localization |
| *C5* | *CPS1* | 0.010474401 | Co-localization |
| *C5* | *ARG1* | 0.010341191 | Co-localization |
| *C5* | *BAAT* | 0.010462028 | Co-localization |
| *G6PC* | *SLC25A15* | 0.016094055 | Co-localization |
| *G6PC* | *HAL* | 0.008763849 | Co-localization |
| *G6PC* | *CPS1* | 0.009082945 | Co-localization |
| *G6PC* | *BAAT* | 0.00880316 | Co-localization |
| *G6PC* | *HGD* | 0.014180019 | Co-localization |
| *G6PC* | *MTHFS* | 0.005369584 | Co-localization |
| *G6PC* | *MBL2* | 0.007431859 | Co-localization |
| *G6PC* | *C5* | 0.011984772 | Co-localization |
| *DIO1* | *SLC25A15* | 0.025056243 | Co-localization |
| *DIO1* | *HGD* | 0.017162617 | Co-localization |
| *LBP* | *SLC25A15* | 0.014860843 | Co-localization |
| *LBP* | *HAL* | 0.008506832 | Co-localization |
| *LBP* | *BAAT* | 0.006969759 | Co-localization |
| *LBP* | *HGD* | 0.011018015 | Co-localization |
| *LBP* | *MTHFS* | 0.004518892 | Co-localization |
| *LBP* | *MBL2* | 0.006218935 | Co-localization |
| *LBP* | *C5* | 0.010104029 | Co-localization |
| *LBP* | *G6PC* | 0.009210799 | Co-localization |
| *SLC17A2* | *SLC25A15* | 0.018329779 | Co-localization |
| *SLC17A2* | *HAL* | 0.009681415 | Co-localization |
| *SLC17A2* | *BAAT* | 0.009294453 | Co-localization |
| *SLC17A2* | *HGD* | 0.013592821 | Co-localization |
| *SLC17A2* | *MTHFS* | 0.006027925 | Co-localization |
| *SLC17A2* | *ADCK2* | 0.015540332 | Co-localization |
| *SLC17A2* | *MBL2* | 0.008607964 | Co-localization |
| *SLC17A2* | *G6PC* | 0.011782652 | Co-localization |
| *SLC17A2* | *LBP* | 0.010201342 | Co-localization |
| *ORM1* | *SLC25A15* | 0.009743731 | Co-localization |
| *ORM1* | *HAL* | 0.005098609 | Co-localization |
| *ORM1* | *CPS1* | 0.005120896 | Co-localization |
| *ORM1* | *ARG1* | 0.004895931 | Co-localization |
| *ORM1* | *BAAT* | 0.004949312 | Co-localization |
| *ORM1* | *C4BPB* | 0.009870878 | Co-localization |
| *ORM1* | *HGD* | 0.007168311 | Co-localization |
| *ORM1* | *MTHFS* | 0.003196832 | Co-localization |
| *ORM1* | *ADCK2* | 0.007305417 | Co-localization |
| *ORM1* | *MBL2* | 0.004552235 | Co-localization |
| *ORM1* | *C5* | 0.006662394 | Co-localization |
| *ORM1* | *G6PC* | 0.006180191 | Co-localization |
| *ORM1* | *LBP* | 0.005398762 | Co-localization |
| *ORM1* | *SLC17A2* | 0.006999721 | Co-localization |
| *SERPINA7* | *SLC25A15* | 0.010623649 | Co-localization |
| *SERPINA7* | *CPS1* | 0.006563142 | Co-localization |
| *SERPINA7* | *ARG1* | 0.006524967 | Co-localization |
| *SERPINA7* | *BAAT* | 0.006416609 | Co-localization |
| *SERPINA7* | *MBL2* | 0.005163964 | Co-localization |
| *SERPINA7* | *C5* | 0.00801424 | Co-localization |
| *SERPINA7* | *G6PC* | 0.007043994 | Co-localization |
| *SERPINA7* | *LBP* | 0.005624746 | Co-localization |
| *SERPINA7* | *SLC17A2* | 0.007645443 | Co-localization |
| *SERPINA7* | *ORM1* | 0.004100054 | Co-localization |
| *CYP3A5* | *SLC25A15* | 0.021976126 | Co-localization |
| *CYP3A5* | *HAL* | 0.012220652 | Co-localization |
| *CYP3A5* | *CPS1* | 0.011446337 | Co-localization |
| *CYP3A5* | *BAAT* | 0.011028161 | Co-localization |
| *CYP3A5* | *MTHFS* | 0.006549534 | Co-localization |
| *CYP3A5* | *C5* | 0.015651088 | Co-localization |
| *CYP3A5* | *G6PC* | 0.01435386 | Co-localization |
| *CYP3A5* | *LBP* | 0.012599541 | Co-localization |
| *CYP3A5* | *SLC17A2* | 0.014993709 | Co-localization |
| *CYP3A5* | *ORM1* | 0.008000645 | Co-localization |
| *CYP3A5* | *SERPINA7* | 0.008649936 | Co-localization |
| *PREP* | *CTH* | 0.047972582 | Physical Interactions |
| *C4BPB* | *SLC25A15* | 0.15971017 | Physical Interactions |

This study is the first in silico analysis to determine the impact of SNPs on the coding region of the SLC25A15, other studies relied on molecular work investigated, specifically the effect of mutations on *SLC25A15* that specified and associated with phenotype of HHH syndrome (26). Also, detailed information on HHH was carried out by analysis of *SLC25A15* gene with computational modeling NMR, in order to study and construct the 3D structure and mutated mitochondrial ornithine transporter 1, and also to provide molecular docking to identify substrate binding site followed by dynamic simulation to optimize these model (51).

20 pathogenic SNPs effect on the SLC25A15 protein function and structure were reveled in this study. 12 of these 20 missense SNPs are proposedly to be considered as novel mutations, while; L71Q, G113C, F188S, G216S, G220R, Q238R, R275Q, and R275L are not originals, due to their previous mentions in studies (1, 11, 12, 26, 52, 53).

NCBI and this study both acknowledge that L71Q, G113S, G220R, and R275Q SNPs to be pathogenic, in addition to, this study also found further 16 other pathogenic SNPs. However underneath NCBI pathogenic results; G27R was to be positive, while in this study G27R resulted as not pathogenic SNP.

Overexpression of *SLC25A15* may enhance the melanoma proliferation, which might affect the melanoma prognosis and future orientation for melanoma therapy (53).

The diagnosis of HHH associates with clinical manifestation of the syndrome, while treatment may improve with protein free diet and administration of arginine, however liver transplant may still be required (54). This new study, it is not only costless, but can rapidly identify the pathogenic SNPs, specifically targeting mutant *SLC25A15* and may aid in biomarker diagnosis of the disease thus allowing safe novel gene therapy to be performed, from the alternative liver transplant.

## 4. Conclusions

In this study 20 pathogenic SNPs (D31Y, Y64C, G66S, L71Q, G86C, G113C, G168E, T176A, F188S, G216S, G217R, G220R, K234I, Q238R, R275Q, R275L, Y287H, E288K, and E288G) were found in SLC25A15 gene through a combination of different in silico bioinformatics tools. Their impact on *SLC25A15* gene functions and structure were also analyzed, thus providing knowledge as a starting point for innovation of new, useful SNP markers for medical testing and a safer medication to treat the most common demoralizing disorders of SLC25A15 in urea cycle.

